# Two-dose “extended priming” immunization amplifies humoral immune responses by synchronizing vaccine delivery with the germinal center response

**DOI:** 10.1101/2023.11.20.563479

**Authors:** Sachin H. Bhagchandani, Leerang Yang, Laura Maiorino, Elana Ben-Akiva, Kristen A. Rodrigues, Anna Romanov, Heikyung Suh, Aereas Aung, Shengwei Wu, Anika Wadhera, Arup K. Chakraborty, Darrell J. Irvine

## Abstract

“Extended priming” immunization regimens that prolong exposure of the immune system to vaccines during the primary immune response have shown promise in enhancing humoral immune responses to a variety of subunit vaccines in preclinical models. We previously showed that escalating-dosing immunization (EDI), where a vaccine is dosed every other day in an increasing pattern over 2 weeks dramatically amplifies humoral immune responses. But such a dosing regimen is impractical for prophylactic vaccines. We hypothesized that simpler dosing regimens might replicate key elements of the immune response triggered by EDI. Here we explored “reduced ED” immunization regimens, assessing the impact of varying the number of injections, dose levels, and dosing intervals during EDI. Using a stabilized HIV Env trimer as a model antigen combined with a potent saponin adjuvant, we found that a two-shot extended-prime regimen consisting of immunization with 20% of a given vaccine dose followed by a second shot with the remaining 80% of the dose 7 days later resulted in increased total GC B cells, 5-10-fold increased frequencies of antigen-specific GC B cells, and 10-fold increases in serum antibody titers compared to single bolus immunization. Computational modeling of the GC response suggested that this enhanced response is mediated by antigen delivered in the second dose being captured more efficiently as immune complexes in follicles, predictions we verified experimentally. Our computational and experimental results also highlight how properly designed reduced ED protocols enhance activation and antigen loading of dendritic cells and activation of T helper cells to amplify humoral responses. These results suggest that a two-shot priming approach can be used to substantially enhance responses to subunit vaccines.

## INTRODUCTION

Vaccines are a critical public health tool for the control of infectious diseases and, most recently, their efficacy in mitigating morbidity and mortality during the COVID-19 pandemic has further highlighted their importance (*1*). However, despite significant advances, there remain a number of pathogens, including the human immunodeficiency virus (HIV), for which effective vaccines are unavailable, owing to challenges such as high mutation rates, immune evasion mechanisms, and unfavorable immunodominance patterns (*2, 3*). HIV continues to pose a persistent global health threat, and the development of an effective vaccine against the virus remains an urgent priority (*4*). Broadly neutralizing antibodies that can neutralize diverse strains have been isolated from naturally infected people (*5*) and they target relatively conserved regions of the HIV envelope spike. Based on non-human primate and human studies of antibody passive transfer, a vaccine capable of eliciting broadly neutralizing antibodies (bnAbs) against the HIV envelope trimer (Env) should be protective against HIV infection (*6–8*). However, generation of antibodies capable of broad and potent native HIV neutralization via vaccination has proven challenging due to multiple factors including immunodominance of non-neutralizing epitopes, high sequence variability of the trimer, and difficult structural accessibility of highly conserved epitopes (*3, 9, 10*). Similar challenges have hindered the development of universal influenza vaccines that target the relatively conserved receptor binding site of hemagglutinin (HA) (*11, 12*).

Following vaccination, germinal centers (GCs) play a pivotal role in the evolution of the clonality and affinity of the antibody response, and influence the composition of the memory B cell and long-lived plasma cell compartments following immunization (*13, 14*). The provision of antigen to GC B cells by follicular dendritic cells (FDCs), which efficiently capture complement- or antibody-decorated antigen, and support from follicular helper T cells (Tfh), which control GC B cell survival, are crucial factors in this process (*15, 16*). Notably, the size of the early GC response correlates with the magnitude of neutralizing antibodies (Abs) generated by immunization with HIV Env trimers in rhesus macaques (*17*). Furthermore, for difficult-to-neutralize pathogens such as HIV, increasing the number of clones entering the GC increases the likelihood that a rare germline B cell that targets a neutralizing epitope could be selected and undergo affinity maturation (*9, 18*).

One effective approach to enhance the GC response is via the use of “extended prime” dosing regimens for vaccine administration (*19–22*). In this approach, a given dose of antigen and adjuvant are provided over a prolonged window of time compared to traditional bolus vaccine injection, through methods such as repeated injections (*19, 20, 22*), implanted drug delivery devices (*19, 20*), or use of slow-release biomaterials (*23–25*). Among these approaches, a simple strategy that elicits profound changes to humoral immunity is escalating-dose immunization (EDI), where a given dose of vaccine is administered as a series of repeated injections over a period of 2 weeks. For stabilized HIV Env trimer immunogens, EDI using seven injections in an exponentially-increasing dosing pattern has been shown to increase the magnitude of the GC and Tfh response, increase the number of B cell clones entering the GC, increase the size of the memory B cell response, increase autologous tier 2 neutralizing antibody titers, and initiate a GC response that can persist for at least 6 months in non-human primates (*19, 20, 22*).

Although this 7-dose extended-prime regimen is highly effective, it is not practical for mass vaccination. Slow-delivery technologies aiming to mimic the effects of extended dosing regimens following a single injection are in development (*23, 26–28*), but it remains to be seen if these approaches can fully replicate the potency of EDI and they must meet a high bar of safety for wide implementation in vaccine development. Our early computational modeling work seeking to understand the mechanisms of EDI immune responses suggested that the key elements driving strong GC responses following ED immunization are the initiation of B cell priming and GC formation by small amounts of antigen early in the dosing course, followed by later (∼1-2 weeks) arrival of larger quantities of antigen that can be captured in immune complexes formed via newly-produced affinity-matured antibody in the lymph node (*19*). This late-arriving antigen is thus deposited on FDCs at high levels, providing a reservoir of antigen to drive the GC response.

Based on these mechanisms, we hypothesized that a “reduced escalating-dose” immunization could be possible, where we simplify the ED dosing regimen to just 2 injections: a first dose to initiate B cell priming, followed by a second shot 1-2 weeks later that would provide antigen for capture on FDCs. Here, we explored the parameter space of EDI and carried out systematic studies varying the number of doses, dose ratio, and dose intervals using a model HIV Env stabilized trimer immunogen and potent saponin adjuvant in mice. We found that a reduced 2-shot extended-prime approach is able to retain much of the benefit of the 7-dose EDI regimen in amplifying the humoral response against Env trimers. Guided by computational modeling of mechanistic steps required for T cell and B cell priming following bolus or multi-shot immunization, we show that even a two-shot priming approach is capable of greatly augmenting antigen deposition on FDCs to drive the GC response compared to a bolus priming immunization. Together these data suggest that a simple extended-prime immunization approach for subunit vaccines could provide substantial enhancements to humoral immunity. Our results also provide new mechanistic insights into how the modulation of vaccine kinetics can be leveraged to augment the germinal center response of broad relevance to vaccines for infectious diseases.

## RESULTS

### A two-dose priming regimen greatly augments responses to HIV Env trimer protein immunization over traditional bolus immunization

Using a stabilized HIV Env SOSIP trimer immunogen engineered to promote priming of N332-supersite-directed B cells (N332-GT2 (*29*)) and a potent saponin nanoparticle adjuvant (saponin/MPLA nanoparticles, SMNP (*30, 31*)), we conducted an evaluation of varied EDI dosing regimens in mice. Given the large parameter space to explore, we opted to focus on analysis of GC responses at day 14 for all groups, as we previously found that GC responses in mice peaked at this timepoint for immunization patterns as disparate as bolus injection and the full 7-dose ED regimen (*19*). First, we tested the effect of number of doses, starting from the previously defined optimal 7-ED regimen, and reducing the number of doses systematically, while keeping the escalation-over-time pattern, the total time interval (12 days), and amount of total vaccine administered (summing all the doses for a given regimen) constant (**Fig. 1A**). As the number of doses was reduced, the total size of the GC and Tfh responses steadily dropped (**Fig. 1B-C**). However, by staining with fluorescent trimer probes to identify antigen-specific cells, we found the number of trimer-binding GC B cells dropped only ∼5-fold moving from 7 doses down to 3 doses, while a 2-dose ED pattern elicited an antigen-specific GC B cell response not statistically different from bolus (**Fig. 1D**). This poor response to the two-dose ED immunization was not due to the choice of time point for GC analysis, as Tfh and antigen-binding GC B cell numbers were also not statistically different from bolus immunization for the 2-dose regimen when measured 7 days after the second dose (**fig. S1A-C**). Trimer-specific serum IgG titers measured one month after dosing were similar for 7, 6, 4, and 3-dose ED regimens, but 2-dose and bolus immunizations elicited antibody titers 8- and 60-fold lower, respectively (**Fig. 1E**).

**Figure 1.**
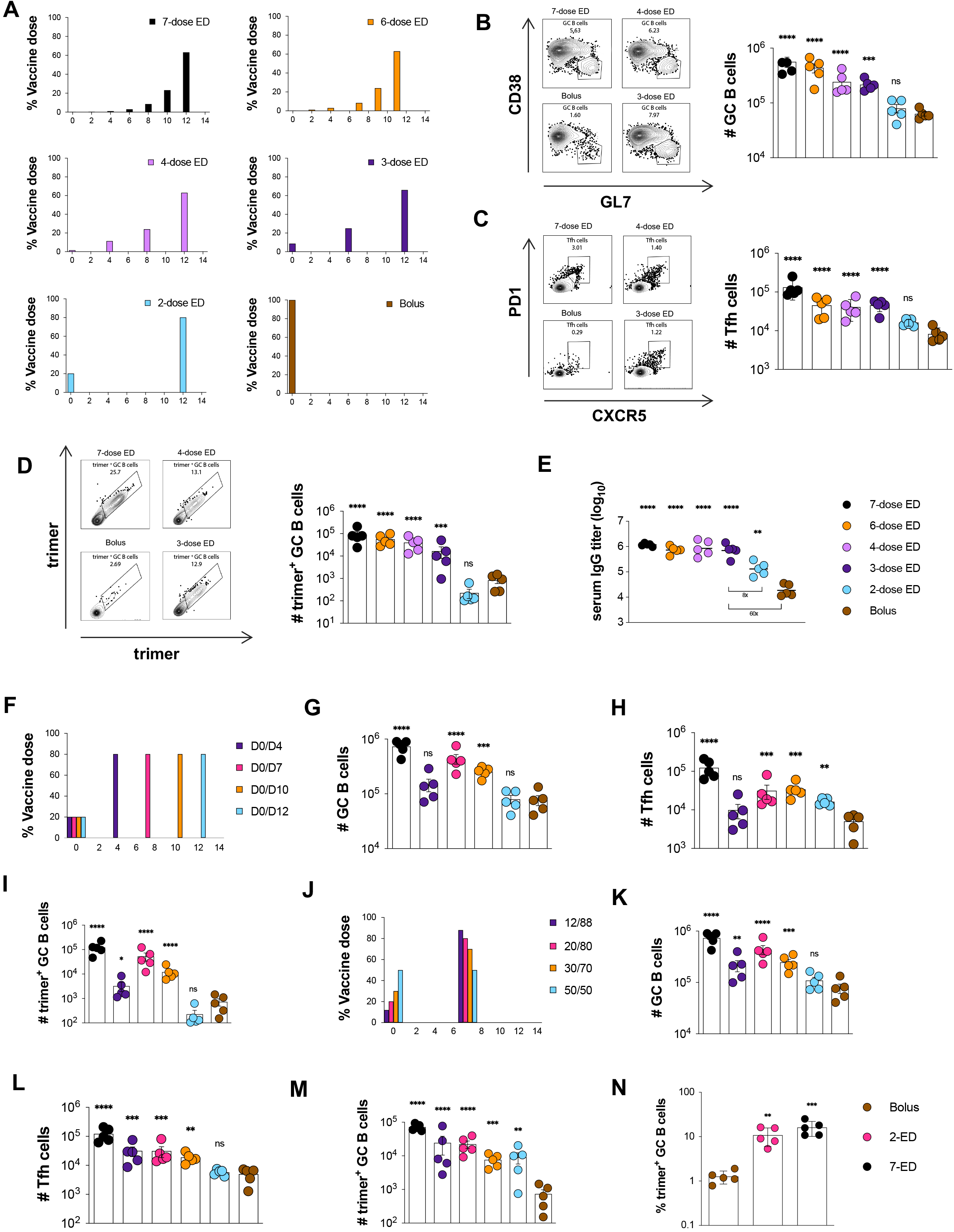
An optimally designed two-shot extended-prime vaccination substantially enhances GC responses to subunit vaccines compared to bolus immunization. **(A)** Schematic of escalating dose vaccination regimens with varying dose number. (**B-E**) C57BL/6J mice (*n*=5 animals/group) were immunized with 10 µg N332-GT2 trimer and 5 µg SMNP adjuvant according to the dosing schemes in (**A**). GC responses were evaluated on day 14 by flow cytometry and antibody responses by ELISA on day 28. Shown are representative flow cytometry histograms and cell counts for total GC B cells (**B**), Tfh (**C**), and trimer-specific GC B cells (**D**) at day 14, and trimer-specific serum IgG titers at day 28 (**E**). (**F**) Schematic of dosing schedules tested for two-shot ED regimens. (**G-I**) C57BL/6J mice (*n*=5 animals/group) were immunized with 10 µg N332-GT2 trimer and 5 µg SMNP adjuvant according to the dosing schemes in (**F**), and total GC B cells (**G**), Tfh cells (**H**), and trimer-specific GC B cells (**I**) were analyzed by flow cytometry on day 14. Note: Bolus and 7-dose ED comparisons are also shown with black and brown colors respectively. (**J**) Schematic of dosing ratios evaluted for for 2-shot ED immunization. (**I-M**) C57BL/6J mice (*n*=5 animals/group) were immunized with 10 µg N332-GT2 trimer and 5 µg SMNP adjuvant according to the dosing schemes in (**J**), and total GC B cells (**K**), Tfh cells (**L**), and trimer-specific GC B cells (**M**) were analyzed by flow cytometry on day 14. (**N**) Frequencies of GC B cells recognizing intact trimer antigen for bolus, optimized 2-ED, and 7-ED regimens. Points represent responses of individual animals while bars indicate mean± s.e.m. Shown are data from one representative of two independent experiments for each immunization series. ****, p < 0.0001; ***, p < 0.001; **, p < 0.01; *, p < 0.05; ns, not significant; by one-way ANOVA with Dunnett’s multiple comparison post test compared to bolus immunization.

We hypothesized that the poor response to 2-dose escalating immunization could be because a 12-day interval between doses is too wide a gap to optimally feed antigen to the GC response, and thus we next tested two-dose escalating patterns administered at intervals ranging from 4 to 12 days, fixing the initial dose at 20% of the total, and the remaining 80% of the vaccine dose administered at the second injection (**Fig. 1F**). In this series, a two-dose escalating prime immunization with an interval of 7 days elicited an optimal response, eliciting 4-fold more total GC B cells and 5-fold more Tfh than bolus immunization (**Fig. 1G-H**). Remarkably, this 7-day 2-dose pattern elicited 10-fold more trimer-binding GC B cells than bolus immunization, only 3-fold fewer than the previously-optimized 7-dose two-week escalating dosing pattern (**Fig. 1I**).

Motivated by these findings, we next evaluated the impact of the dosing ratio (the proportion of total vaccine administered at dose 1 vs. dose 2, **Fig. 1J**). Administration of 20% of the vaccine in the first dose was optimal, while increasing the initial immunization to 30 or 50% of the total dose led to decaying responses (**Fig. 1K-M**). Notably, this optimized 2-shot escalating dose regimen administering 20% of the vaccine at day 0 and 80% at day 7 (hereafter, 2-ED) induced similar responses as administration of 2 full vaccine doses a week apart, suggesting that additional antigen at the first dose cannot further augment the GC response (**fig. S1D-H**). The enhanced elicitation of antigen-binding GC B cells elicited by two-dose ED priming reflects a combination of increased size of the GC as well as an increased proportion of GC B cells recognizing intact antigen– with 2-ED dosing eliciting a 6-fold greater frequency of trimer-binding GC B cells compared to bolus vaccination (**Fig. 1N**). Altogether, these data demonstrate that even a minimal two-dose extended priming immunization augments multiple facets of the humoral response to vaccination.

### Optimized 2-dose ED priming amplifies the magnitude but does not alter overall lifetime of the GC response compared to bolus immunization

We next assessed the evolution of the B cell response over time for the optimized 2-ED regimen compared to bolus immunization to evaluate potential differences in the temporal dynamics or lifetime of the GC in each case (**Fig. 2A**). Trimer-binding B cells were detectable in both groups at day 7 and peaked at day 14, but total antigen-binding B cells were 5-fold greater in the 2-ED group compared to bolus (**Fig. 2B**). Interestingly, despite the difference in initial vaccine dose administered, GC responses for bolus and 2-dose ED were similar at day 7, but diverged at day 14 (**Fig. 2C**). GCs in both groups then contracted from day 14 through day 28. The number of Tfh cells in the 2-ED group also sharply expanded between day 7 and 14, reaching ∼6-fold greater numbers of Tfh cells at the peak of the response at day 14 compared to bolus immunization (**Fig. 2D**). Trimer-binding GC B cell numbers also peaked for both immunization conditions at day 14, and the 2-ED regimen maintained higher levels of antigen-specific GC cells compared to bolus immunization through day 21 (**Fig. 2E**). These data demonstrate that a two-dose escalating prime immunization alters the magnitude but not the lifetime of the GC response.

**Figure 2.**
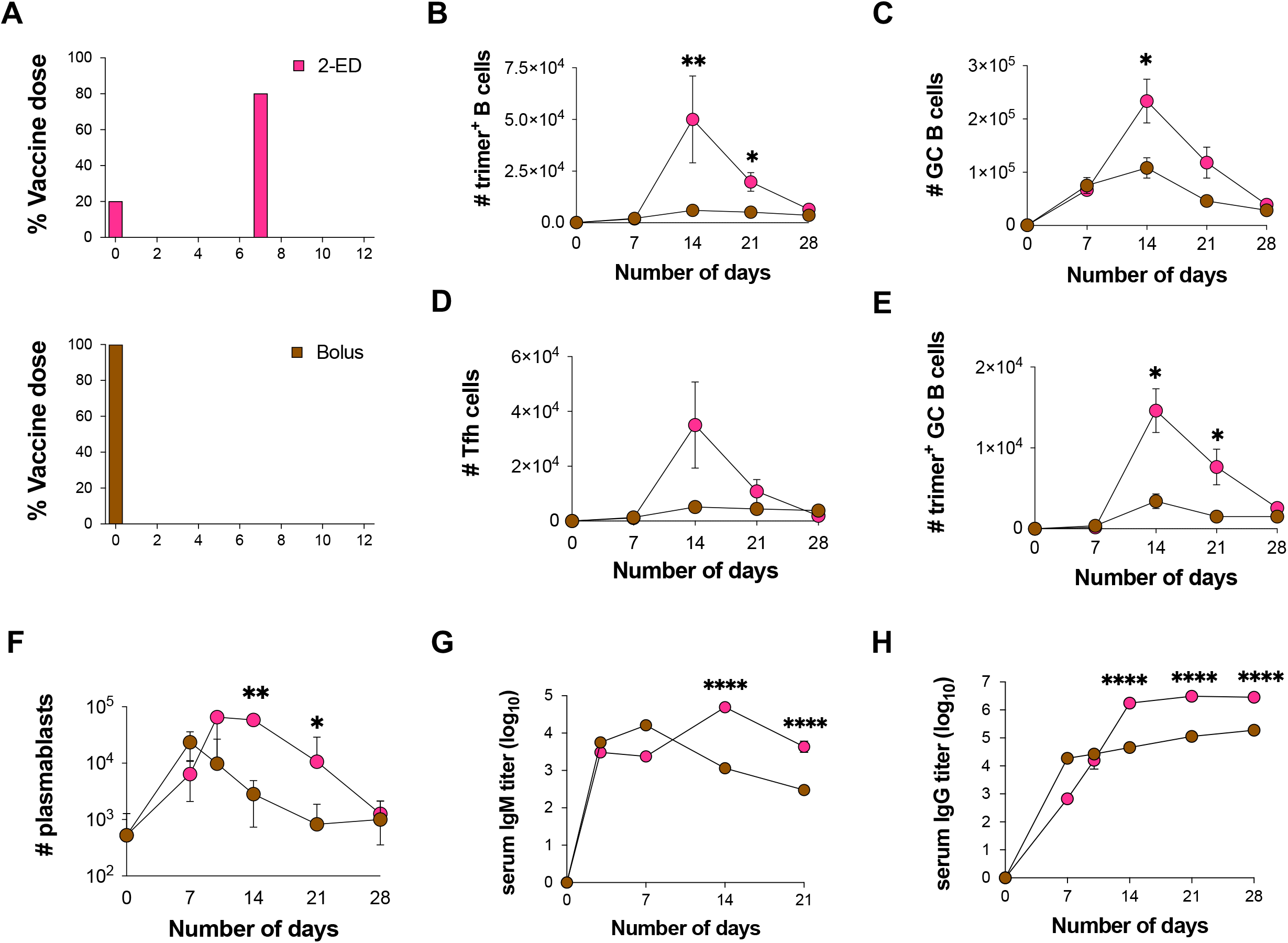
Optimized 2-shot prime immunization amplifies the GC response and trimer-specific serum antibody titers over time compared to bolus immunization. (**A**) Schematic of dosing schemes. (**B-H**) C57BL/6J mice (*n*=9 animals/group) were immunized with 10 µg N332-GT2 trimer and 5 µg SMNP adjuvant according to the dosing schemes in (**A**). GC responses were evaluated on days 7, 14, 21, and 28 by flow cytometry and antibody responses by ELISA on days 3, 7, 14, 21, and 28. Shown are trimer-specific B cell counts (**B**), GC B cell counts (**C**), Tfh cell counts (**D**), trimer-specific GC B cell counts (**E**), plasmablast counts (**F**), trimer-specific IgM titers (**G**), and trimer-specifc IgG titers (**H**), plotted over time for bolus and 2-ED regimens. Shown are data from one independent experiment for each immunization series. ****, p < 0.0001; ***, p < 0.001; **, p < 0.01; *, p < 0.05; ns, not significant; by two-way ANOVA with Dunnett’s multiple comparison post test compared to bolus immunization.

Lymph node plasmablasts were expanded by day 7 for both the bolus and 2-ED immunizations; these responses peaked at day 7 and day 10, respectively, then steadily decayed (**Fig. 2F**). ED immunizations also impacted the evolution of serum antibody responses. IgM responses primed by bolus immunization peaked at day 7 then decayed steadily, while the 2-ED group elicited IgM responses that peaked later, at day 14 (**Fig. 2G**). Bolus immunization elicited substantial serum IgG titers by day 7 which rose only slightly over the subsequent 3 weeks (**Fig. 2F**). By contrast, IgG responses for the 2-ED regimen increased sharply between day 7 and day 14, reaching levels ∼10-fold greater than the bolus group by day 14 that were maintained over time (**Fig. 2H**). Hence, the simplified 2-dose ED regimen elicited changes in the humoral response that persisted over many weeks and primed strong, stable serum antibody responses that were greatly increased compared to traditional bolus immunization.

### Extended-prime dosing regimens boost innate inflammation in lymph nodes and allow for improved T cell responses

We next sought to understand the mechanisms underlying the substantial amplification of the humoral immune response obtained by 2-ED prime relative to traditional bolus immunization. A strong correlation between the number of Tfh and GC B cells has been observed in prior studies (*32, 33*), and is consistent with selection by Tfh cells serving as a key bottleneck controlling proliferation of GC B cells (*13, 14, 34, 35*). We hypothesized that the relative kinetics of antigen and adjuvant availability during ED vs. bolus immunization could substantially impact steps in the priming of Tfh cells, and that this may be one important mechanism governing responses to 2-ED dosing. To test this idea, we developed a coarse-grained kinetic computational model of the innate immune response and T cell priming following vaccine administration (**Fig. 3A, fig. S2A**). Briefly, in this simple model, both antigen (Ag) and adjuvant (Adj) appear as a bolus at an initial administered concentration and clear from the tissue at a constant rate. The adjuvant recruits and activates tissue cells at the immunization site and/or draining lymph node (**Fig. 3A, (i)**), which release cytokines and chemokines that recruit dendritic cells (DCs) (**Fig. 3A, (ii)**). Adjuvant also plays an important role in activating DCs and promoting antigen uptake by these cells (*36*). As a proxy for these effects, we assume that the adjuvant increases the rate of antigen uptake by DCs and induces DC activation in an Adj and Ag concentration-dependent manner (**Fig. 3A, (iii)**). Activated, antigen-loaded DCs (aDC^Ag+^s) then prime T cells in the draining lymph node (**Fig. 3A, (iv)**). The proliferation of T cells induced by the aDC^Ag+^s is described according to a previously reported model (*37*), and we assume proliferating T cells differentiate into Tfh cells at a constant rate (**Fig. 3A, (v)**). This minimal model is easily interpretable and has a small number of parameters, most of which can be estimated from previous experimental studies (See **Supplementary Materials** and **Table S2** for details on the model and parameter estimation) (*38*).

**Figure 3.**
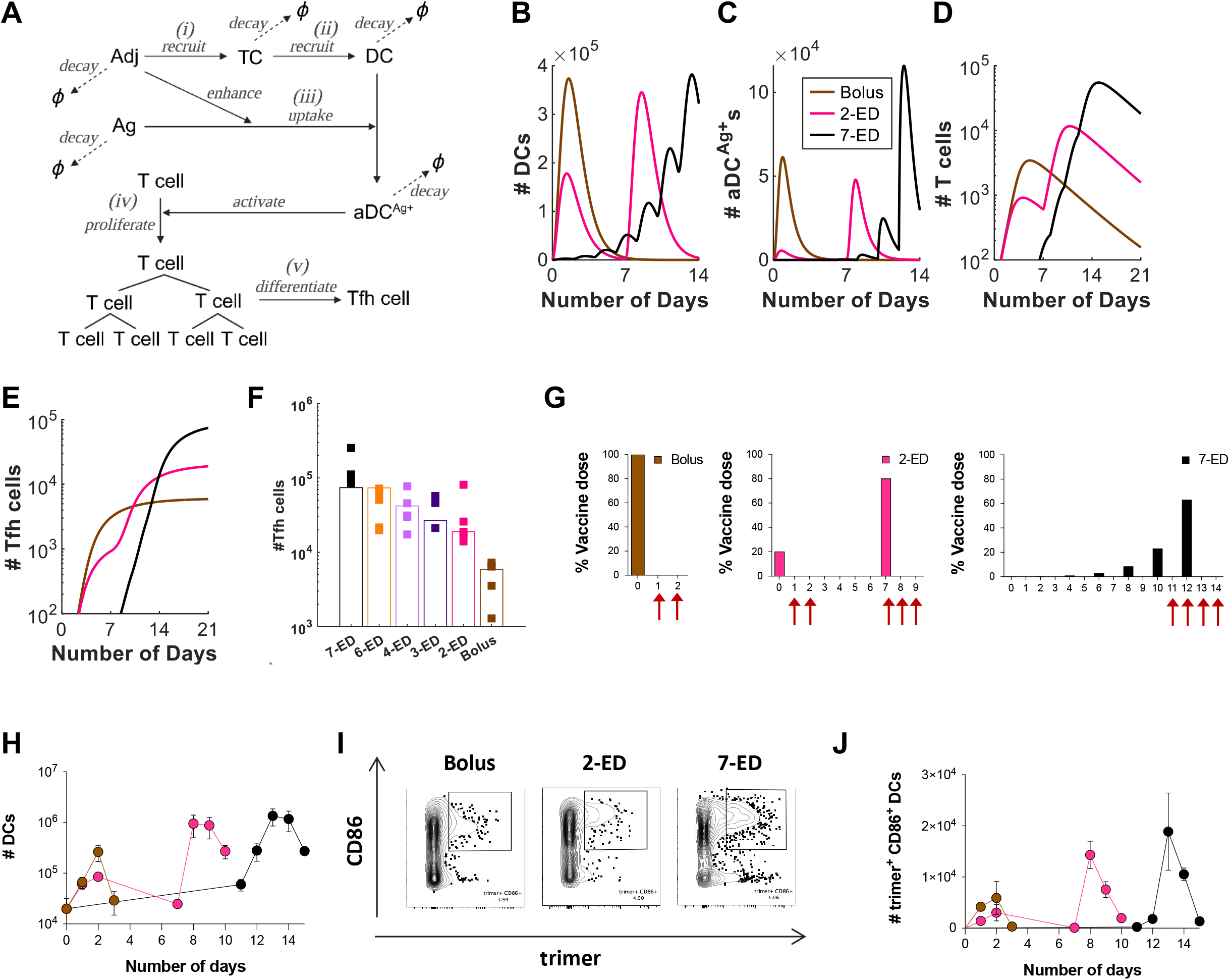
In Silico modeling predicts enhanced T cell priming with extended-prime immunization, consistent with experimental measurements of DC antigen acquisition and activation in draining lymph nodes. (**A-F**) Computational model of vaccine uptake by dendritic cells and helper T cell priming. (**A**) Schematic outlining elements of the kinetic model. (**B-E**) Modeling predictions of the number of (**B**) total DCs (**B**), Ag^+^Adj^+^ DCs (**C**), Ag-specific T cells (**D**), and Tfh cells (**E**) for bolus, 2-ED, or 7-ED immunization regimens. (**F**) Comparing Tfh cell count predicted by the model with the experimental data at day 14. (**G-J**) Experimental analysis of lymph node DC antigen uptake and activation. C57BL/6J mice (*n*=3 animals/group) were immunized with 10 µg Cy5 dye-labled-N332-GT2 trimer and 5 µg SMNP adjuvant according to the dosing schemes shown in (**G),** and DCs in draining lymph nodes were analyzed by flow cytometry on days indicated by arrows. Shown are number of DCs (**H**), representative histograms of trimer antigen fluorescence and CD86 expression by CD11c^+^ DCs (**I**), and number of trimer^+^CD86^+^ DC counts over time for bolus, 2-ED, and 7-ED immunization regimens (**J**).

We first examined predictions of the number of total DCs, aDC^Ag+^s, T cells, and Tfh cells following bolus, optimal 2-ED, and 7-ED immunization schemes (**Fig. 3B-E**). After bolus immunization, both total DCs and aDC^Ag+^s reach their peak within ∼1 day and then decline due to rapid adjuvant and antigen clearance (**Fig. 3B-C**). In contrast, the 7-ED regimen shows DCs being periodically recruited with each dose, leading to accumulation in lymph nodes from day ∼7 onward (**Fig. 3B**). The number of aDC^Ag+^s also gradually increases, reaching a large peak value after the final dose on day 12 (**Fig. 3C**). Given the exponential nature of T cell proliferation, extended stimulation by the increasing numbers of aDC^Ag+^s over time during the 7-ED regimen contributes to the substantially greater peak T cell and Tfh cell counts compared to the bolus (**Fig. 3D-E**), consistent with previous studies (*37, 39*). The model suggests that the 7-ED regimen also benefits from efficient generation of aDC^Ag+^. Unlike bolus immunization, where antigen decays before most DCs are recruited, a substantial number of DCs are already present in the lymph node when the final dose of 7-ED regimen is given on day 12. This leads to a predicted prominent increase in the number of aDC^Ag+^s (**Fig. 3B-C**). For the 2-ED regimen, the model predicts T cells proliferate after the 1^st^ dose, and remain at elevated numbers at the time of the second dose (**Fig. 3D**). This expanded pool of antigen-specific T cells will then continue to expand on stimulation with the second dose of vaccine. However, with realistic biological parameters, this simple model predicts that DC numbers elevated after the first dose returns to baseline before the second dose (**Fig. 3B**).

We next modeled each of the escalating-dose regimens tested in **Fig. 1C**, and found that the model indeed captured the pattern of Tfh responses observed experimentally, including the enhanced Tfh response elicited by a day 0/day 7 two-dose vaccination (**Fig. 3F**). Note that two parameters in the model, those corresponding to the rate of DC recruitment and the baseline number of T cells, were fitted directly to the experimental data in **Fig. 3F** (See **Supplementary Material** for discussion on these parameters). Because of the relationship between adjuvant and DC recruitment/activation, the model also predicts that optimal ED immunization requires co-administration of antigen and adjuvant across the time course-if antigen is administered in an escalating-dose pattern but adjuvant is administered as a bolus, poor synchronization of DC recruitment and antigen availability is predicted, and thus weak Tfh priming despite escalating dose antigen administration (**fig. S2B-C**). In agreement with this prediction, vaccination of mice with a bolus adjuvant/7-ED antigen dosing schedule elicited much weaker GC and Tfh responses than escalating dosing of antigen and adjuvant together (**fig. S2D-E**). Thus, this very simple model captures the relative magnitudes of increased Tfh expansion observed for the two ED regimens compared to traditional bolus immunization.

To test the predictions of the kinetic model, we immunized mice with fluorescently-labeled N332-GT2 trimer and SMNP adjuvant, using ED and bolus dosing schemes. We then analyzed DCs in the draining lymph nodes (dLNs) at key time points post immunization (**Fig. 3G, fig. S3**). After bolus immunization, both the total number of DCs and activated (CD86^+^) trimer^+^ DCs in dLNs increased over 2 days (**Fig. 3H-J**). The initial dose of 2-ED similarly expanded DCs and activated antigen-loaded DCs over the first two days, though reaching lower peak numbers. DC numbers in the dLNs returned to baseline by day 7, before the second dose was administered, as predicted by the model (**Fig. 3H, J**). Intriguingly, following the second dose in the 2-ED regimen, the kinetics of DC influx were accelerated compared to the response after the first dose, with a significant increase in activated, antigen-loaded DCs observed as early as 24h post injection (**Fig. 3I, J**). We hypothesize that the faster kinetics of DC recruitment could be a consequence of increased antigen presentation and residual inflammation in the dLN from the first dose. In the 7-ED regimen, the accumulation of trimer^+^ CD86^+^ DCs in the lymph nodes becomes markedly evident around day 12. The buildup of DCs peaked following the last dose of the 7-ED regimen on day 12, with peak trimer^+^CD86^+^ DCs on day 13 more than double those observed with the bolus immunization, in line with model predictions (**Fig. 3I-J**, also compare **Figs. 3J** and **3C**). We note that while the accumulation of antigen-loaded DCs is not substantially measurable before day 12, even a small number of antigen-loaded DCs may initiate T cell proliferation from the baseline, as suggested by the model (**Fig. 3C-D**). Given the exponential nature of T cell proliferation, such early proliferation can have considerable importance.

### Computational modeling of the GC response predicts improved native antigen capture following extended-prime immunizations

Using a computational model of the germinal center reaction, we previously predicted that ED immunization can increase the size of the GC response via capture of antigen on follicular dendritic cells (FDCs) (*19*): initial vaccine doses in the ED regimen trigger initial B cell priming, and 7-10 days after the start of dosing, affinity-matured antigen-specific antibodies will begin to be produced. During ED immunization, antigen is still being delivered to the lymph node at this time point, and these newly produced antibodies form immune complexes with incoming antigen and facilitate its transport to FDCs, where it can promote expansion of the GC response (*19*). However, in the present studies we found that ED dosing also dramatically enhances the proportion of GC B cells that bind to the intact immunogen (10-fold and 6-fold vs. bolus for 7-dose ED and 2-dose ED, respectively, **Fig. 1N**). In other work, we recently showed that extracellular protease activity in lymph nodes can play an important role in modifying B cell responses, as extracellular antigen accumulated in lymph nodes following bolus vaccination undergoes rapid proteolytic degradation over a period of a few days (*40*). This antigen breakdown occurring in sinuses and extrafollicular regions of the node limits the quantity of intact antigen available to B cells and creates immunogen breakdown products that can prime competitive off-target B cell responses. However, protease activity was found to be low in B cell follicles, and antigen captured by FDCs during ED immunization led to a much greater accumulation of intact antigen in follicles for escalating dose vs. traditional bolus immunization (*40*).

To determine if greater levels of intact antigen captured in follicles could explain the greatly increased proportion of trimer-specific GC B cells detected for ED immunization, we modified our computational model of the B cell and antibody response (*35*) (which combines the cellular dynamics of B cell proliferation and GC B affinity maturation with the kinetics of antibody production and antigen capture), to incorporate effects of antigen degradation (see **Supplemental Methods**). Briefly, we assume that B cells can target either native antigen or non-native partially degraded antigen, as schematically shown in **Fig. 4A**. Both types of antigen can be transported to FDCs by the corresponding antibodies that bind to them, where they are protected from further degradation. Additionally, to model the antibody response to HIV Env immunogens, we assume that the non-native antigen is more immunodominant– as HIV Env trimers are heavily glycosylated and present few sites for high affinity antibody binding in the intact state, but are expected to expose more proteinaceous surfaces as they degrade (*9*).

**Figure 4.**
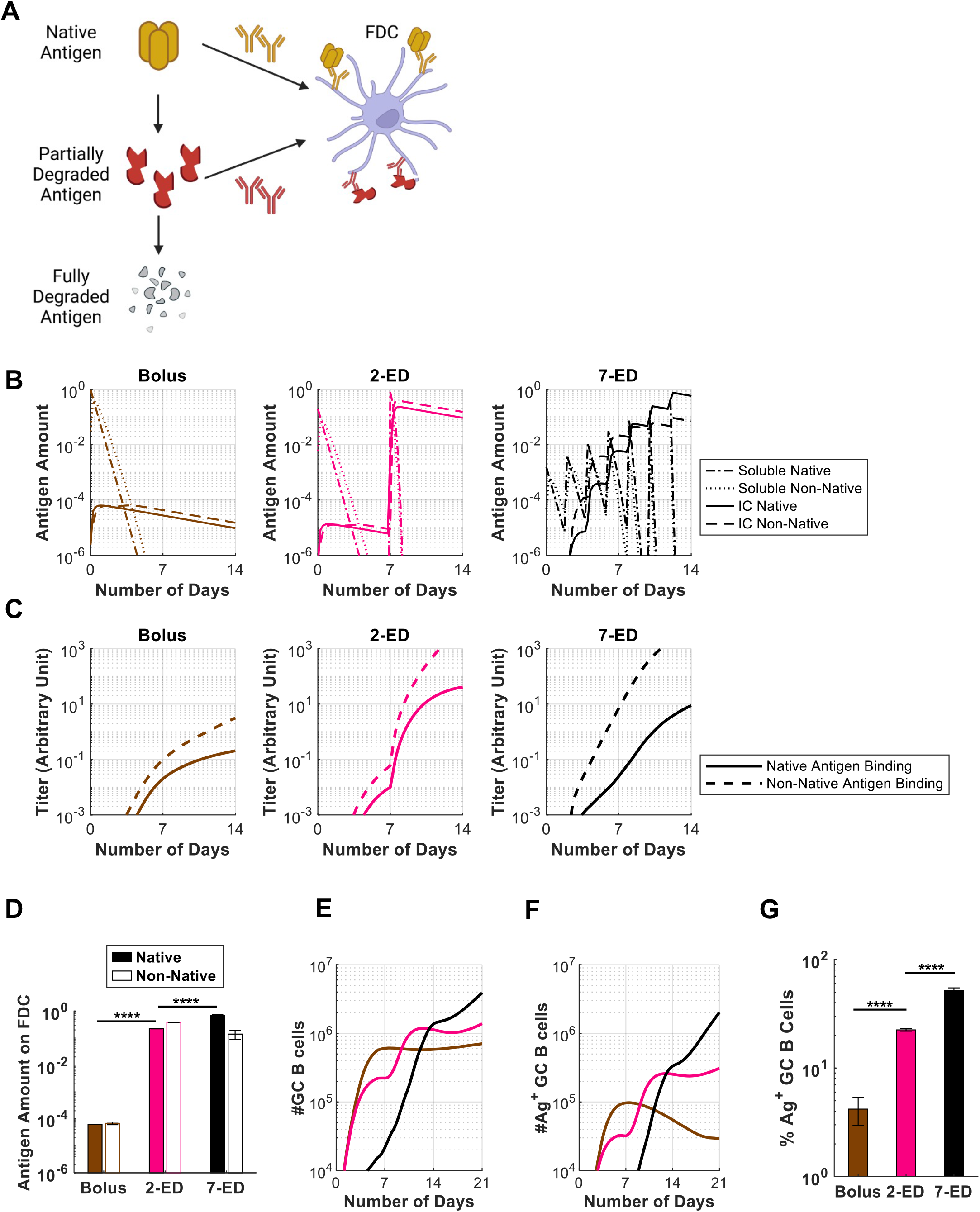
*In Silico* modeling predicts increased intact antigen accumulation on FDCs for extended dosing compared to bolus immunization. (**A**) Schematic showing antigen fates considered in the computational model. (**B**) *In silico* prediction of the levels of free antigen in lymph nodes over time in an intact (“soluble native”) or degraded (“soluble non-native”) state, and amounts of native and non-native antigen captured on FDCs in the form of immune complexes (“IC”) over time. The antigen amounts are normalized to the total antigen dose in immunization. (**C**) Antibody titers predicted by the *in silico* model for bolus, 2-ED, and 7-ED immunization regimens. In the simulation, antibody titers are defined as the concentration of antibodies weighted by their affinities, reflecting their capabilities to bind to the antigen. (**D**) Comparison of predicted antigen amounts accumulated on FDCs after the final shot from each dosing scheme, normalized to the total antigen dose in immunization. (**E**) Model prediction for the number of GC B cells over time. (**F, G**) Model prediction for the number of native antigen-binding (i.e. trimer^+^) GC B cells over time (**F**) and frequency of trimer^+^ GC B cells at day 21 (**G**) from bolus, 2-ED, and 7-ED immunization schemes. The results reported are mean values from 10 independent stochastic simulations of the lymph node. ****, p < 0.0001; by one-way ANOVA with Tukey’s multiple comparison post test.

This revised GC model shows that very small overall amounts of antigen are captured by FDCs following bolus immunization because most of the antigen decays before antigen-specific antibody responses are induced, and very low levels of immune complexes (IC) are formed (**Fig. 4B-D**). As a result, highly stringent conditions are maintained for GC B cell survival, resulting in a GC response that is dominated by the immunodominant non-native antigen-targeting B cells, and the frequency of B cells able to bind to native antigen (Ag^+^ GC B cells) is very low (**Fig. 4E-G**). With the 2-ED regimen, a weak antibody response against both the native and non-native antigens is generated after the first dose (**Fig. 4C**). However, upon the second dose, this modest level of antibody is predicted to efficiently form ICs with incoming antigen leading to a dramatic increase in the amount of antigen deposited on FDCs (**Fig. 4B,D**). The increased antigen availability weakens the immunodominance hierarchy between the two antigens by allowing initially low-affinity B cells to become activated, proliferate and increase affinities through mutations (*35*). This change leads to a much greater native antigen-binding GC B cell response that expands post-second-dose (**Fig. 4E-G**). Finally, with 7-ED dosing, the antibody titer is already high when the final dose (63% of vaccine dose) is administered on day 12 (**Fig. 4C**). Immune complex formation as native antigen arrives at the lymph node at the end of the 7-ED dosing leads to rapid transport of the immunogen to follicles. This causes the ratio of native vs. non-native antigen presented on FDC to be modestly shifted in favor of the former (**Fig. 4D**), allowing even better native antigen-binding responses compared to the 2-dose scheme (**Fig. 4E-G**). Note that native antigen-binding antibody titer at the time of final dose for 7-ED (on day 12) is greater than that for 2-ED at the time of the final dose (on day 7) (**Fig. 4C**). These predictions of the GC model align well with the experimentally-measured frequencies of intact trimer-binding GC B cells at day 14 (compare **Fig. 4G** and **Fig. 1N**).

### A two-dose escalating prime increases antigen capture in follicles compared to bolus immunization

A key prediction of the GC computational model is that a sufficient quantity of antibody specific for intact antigen can be produced by day 7 to enable substantial antigen capture on FDCs using the 2-dose escalating prime immunization. As shown in **Fig. 2G-H**, antigen-specific IgM and IgG were both detectable in serum by day 7 in the 2-ED regimen. To test the model prediction, we immunized mice with fluorescently-labeled N332-GT2 HIV Env trimer by each of the three dosing schedules, and analyzed the biodistribution of antigen in draining lymph nodes via confocal imaging of histological sections and cleared whole lymph node tissues. As seen in prior studies (*19*, *20*), both cleared whole dLNs (**Fig. 5A, fig. S4A-C**) and traditional thin section imaging (**Fig. 5B, fig. S4D**) revealed the presence of substantial amounts of antigen co-localized with FDCs 2 days following the last dose of the full 7-ED regimen, while little if any antigen could be detected at any location within dLNs 2 days after bolus vaccination. Strikingly, substantial amounts of FDC-localized antigen were also found in draining lymph nodes two days following the second dose of the 2-dose ED immunization (**Fig. 5A-B, fig. S4**). High magnification imaging of the follicles of the ED groups suggested this follicle-localized antigen was associated with FDC dendrites (**Fig. 5B**).

**Figure 5.**
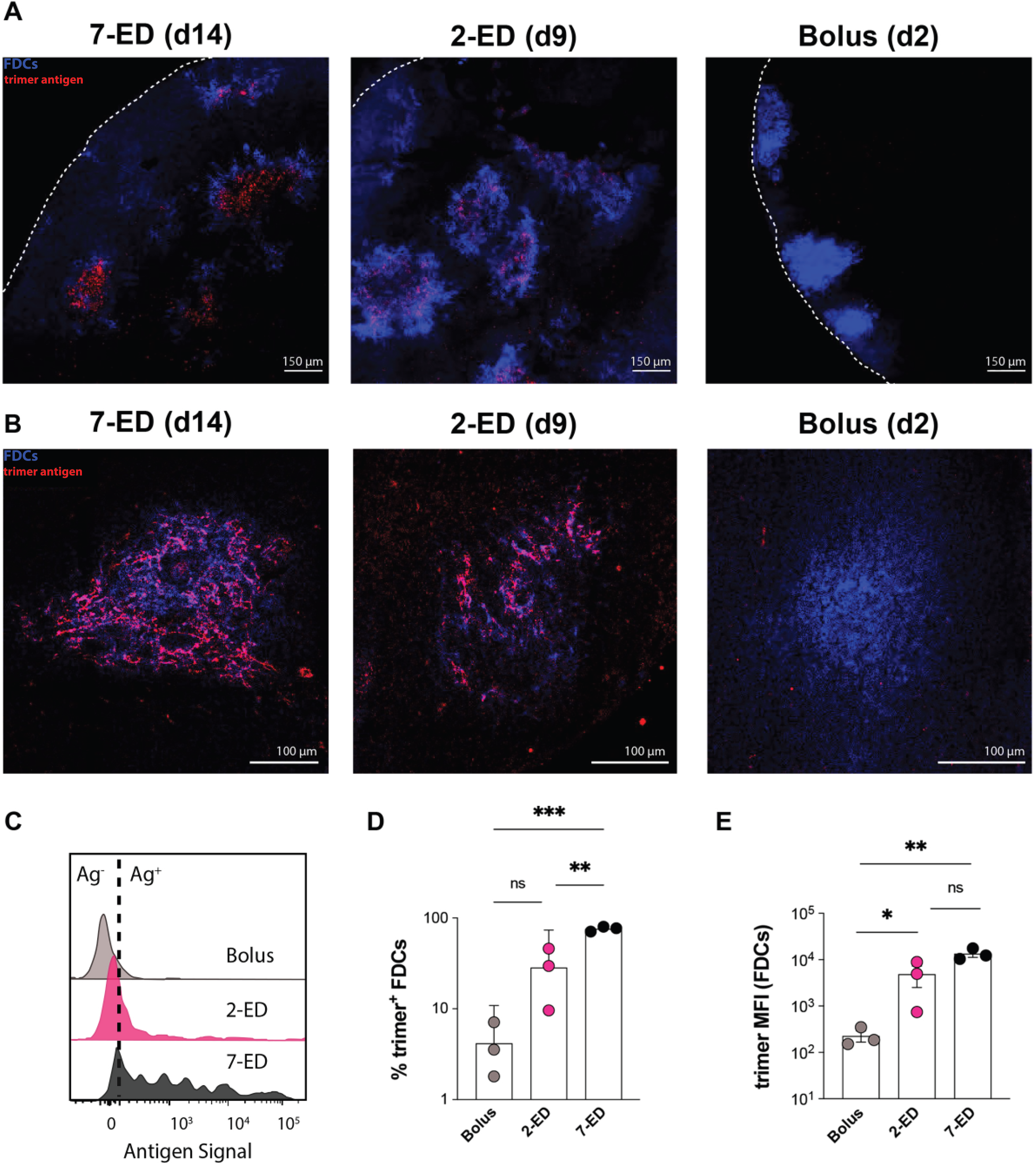
Two-dose extended prime immunization enables antigen capture of the second shot on FDCs. (**A-B**) Groups of C57Bl/6 mice (n=3 animals/group) were immunized by bolus, 2-ED, or 7-ED regimens as in Fig. 3G followed by collection of lymph nodes for imaging at 48 h after bolus or after the last injection of 2-ED and 7-ED regimens. FDC networks were labeled *in situ* by s.c. injection of anti-CD35 antibody 16h before tissue collection. Collected tissues were clarified and imaged intact by confocal microscopy; shown are maximum intensity projections from z-stacks through FDC clusters (Scale bars, 150 μm), (**A**). Alternatively, lymph node sections were stained for FDCs (CD35; blue) and then analyzed by confocal microscopy (Scale bars, 300 μm) to detect co-localization with Cy5-labeled N332-GT2 (pink), (**B**). (**C-E**) Flow cytometry analysis of LN cells (*n*=3 pools/group, with each pool containing six LNs from 3 mice) isolated 48 hr after the final injection following immunization with fluorescently labeled N332-GT2 (10 ug) and SMNP (5 ug) using either bolus, 2-ED, or 7-ED dosing regimens. Shown are representative histograms of antigen intensities among LN cells (**C**), frequencies of trimer+ FDCs (**D**), and the mean trimer fluorescence intensity among trimer^+^ FDCs (**E**) for the indicated immunization conditions. Shown are data from one independent experiment for each immunization series. ****, p < 0.0001; ***, p < 0.001; **, p < 0.01; *, p < 0.05; ns, not significant; by one-way ANOVA with Dunnett’s multiple comparison post test compared to bolus immunization.

Flow cytometry analysis of FDCs recovered from pooled LNs of multiple mice immunized with the different dosing regimens revealed that 2-ED or 7-ED immunizations increased the amount of FDC- trapped antigen by 20-fold and 60-fold over bolus immunization, respectively (**Fig. 5C-E**). Thus, consistent with the computational model of GC dynamics, even a two-dose escalating prime vaccination is capable of achieving antigen targeting to FDCs and greatly increasing the level of antigen retained within the LN.

### Extending antigen availability on the second immunization further boosts humoral responses as a consequence of enhanced native antigen presentation

Given that the computational model of the GC reaction is able to provide qualitatively accurate predictions of the observed GC size and proportion of antigen-specific GC B cells, we next used the model to explore a much wider parameter space of dosing patterns than would be possible experimentally, to gain further insight into how humoral responses might be further bolstered using extended priming. The modeling and experimental data suggest that having a high trimer-binding antibody titer present before the majority of the antigen dose is administered is an important factor governing the magnitude of the “on target” GC response. In the 2-dose extended prime immunization, antibody titers increase steadily after the 2^nd^ injection (**Fig. 2H** and **Fig. 4C**). However, because intact antigen in the lymph node decays rapidly (estimated half-life ∼6.2 hours, **Table S1**), most of the antigen arriving after the second immunization decays before high titers of antibody are reached, leading to presentation of significant amount of non-native antigen on FDCs. Thus, we hypothesized that extending antigen availability over a longer time period at the second dose might further substantially enhance trimer-binding GC responses. We thus simulated a 2-dose ED regimen where antigen was released from the injection site at a constant rate over 10 days after the second injection (**Fig. 6A**). In this case, a significant fraction of the antigen dose arrives at the dLN after high titer levels of trimer-specific antibody are generated and is thus captured in immune complexes in an intact state (**Fig. 6B-C**). Compared to 2-ED dosing using bolus injections, the fraction of intact antigen among ICs increases from ∼38% to 92% with the slow-release second dose (**Fig. 6D**). Thus, the model predicted that such a scheme may lead to a superior trimer-specific GC B cell response compared to the 2-dose immunization (**Fig. 6E-F**). Varying the duration of antigen delivery after the second dose shows that a longer release duration leads to higher fraction of native antigen on FDCs (**Fig. 6G**) and better trimer-specific GC B cell response (**Fig. 6H-I**), highlighting the importance of dosing kinetics.

**Figure 6.**
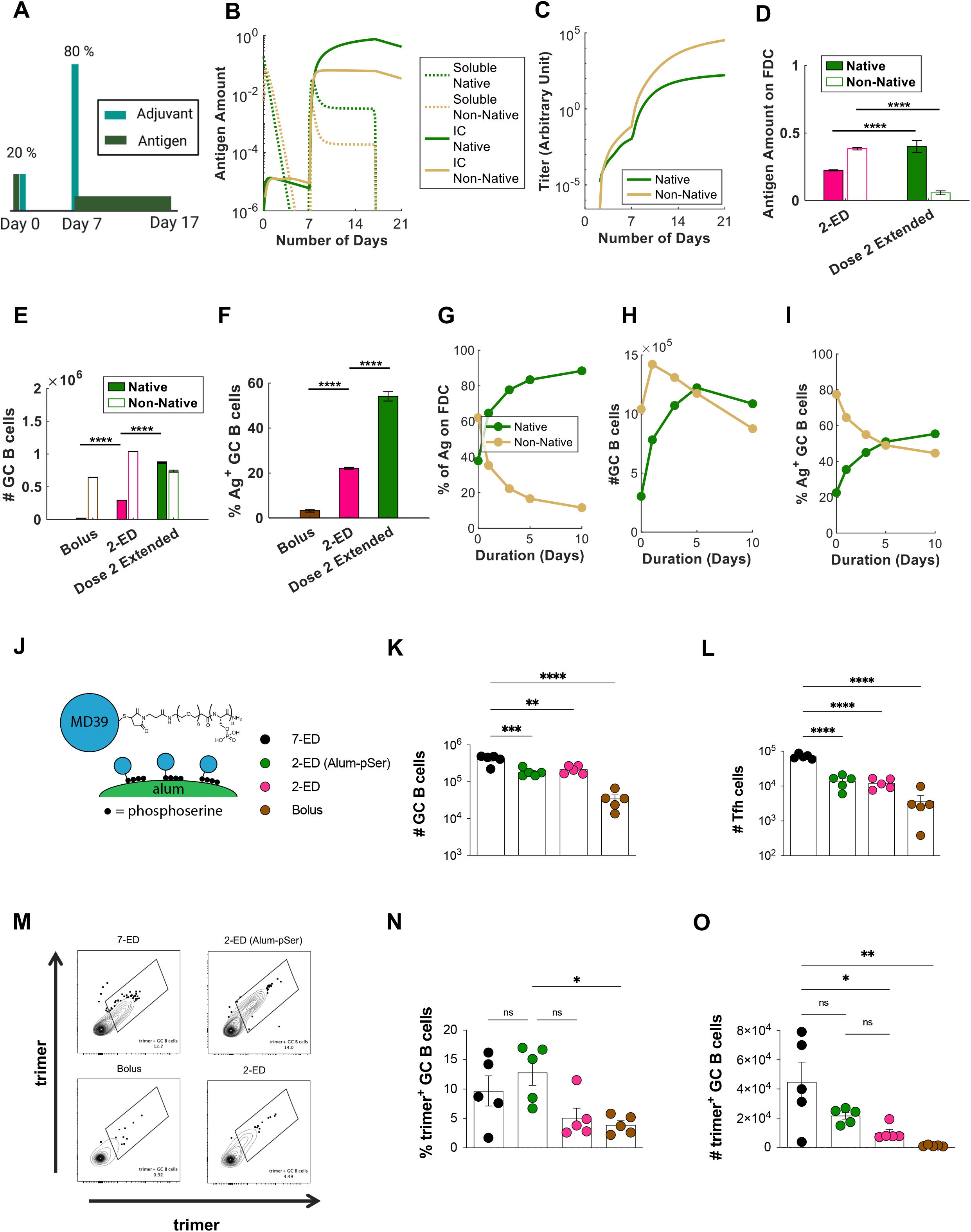
Extending the duration of antigen delivery during the second dose of 2-ED vaccination increases native antigen capture on FDCs and antigen-specific GC responses. (**A-I**) Computational modeling of GC responses elicited by 2-ED dosing administered as two bolus doses vs. a bolus on day 0 and a prolonged antigen delivery at day 7 (“dose 2 extended”). (**A**) Schematic of 2-ED vs. “dose 2 extended” vaccination regimens. (**B, C**) Amounts of free and immune-complexed native or degraded (“non-native”) antigen in the LN over time (**B**) and serum antibody titers recognizing native vs. non-native antigen (**C**) for the “dose 2 extended” regimen. (**D-F**) *In silico* prediction of antigen captured on FDCs (**D**), total GC B cells at day 21 (**E**), and frequency of trimer-binding GC B cells at day 21 (**F**) for bolus, 2-ED, and “dose 2 extended” vaccination regimens. (**G-I**) *In silico* prediction of proportions of intact vs. degraded antigen captured by FDCs (**G**), total number of GC B cells (**H**), and the fraction of GC B cells that are native vs. non-native antigen-binding (**I**) at day 21 as a function of the duration of antigen release used in “dose 2 extended” vaccination. (**J-O**) Experimental testing of “dose 2 extended” immunizations using alum-anchored immunogens. (**J**) Schematic demonstrating anchoring trimer immunogen onto alum via phosphoserine linkers (Alum-pSer). (**K-O**) C57Bl/6J mice (*n*=5 animals/group) were immunized with 10 µg MD39 trimer (either soluble bound to 50 µg alum) and 5 µg SMNP adjuvant as in Fig. 3G. Shown are the numbers of GC B cells (**K**) and Tfh cells (**L**), representative histograms of trimer staining of GC B cells (**M**), frequencies of trimer-binding GC B cells (**N**), and the number of trimer-binding GC B cells (**O**), for the different dosing regimens determined by flow cytometry at day 14. Shown are data from one representative of two independent experiments for each immunization series. ****, p < 0.0001; ***, p < 0.001; **, p < 0.01; *, p < 0.05; ns, not significant; by one-way ANOVA with Dunnett’s multiple comparison post test compared to bolus immunization.

To experimentally test this idea, we employed an approach we previously developed (*27*) to achieve slow-delivery effects in a manner readily translatable to clinical use: a stabilized HIV Env trimer termed MD39 (*41*) (very similar in sequence to the N332-GT2 trimer) was conjugated with short phosphoserine (pSer) peptide affinity tags (one tag per protomer at the C-termini located at the base of the trimer). Phosphate groups in the affinity tags undergo a ligand exchange reaction with the surface of alum particles, enabling oriented high-avidity binding of the trimer to aluminum hydroxide adjuvant (**Fig. 6J**). By stably anchoring the antigen to alum particles, on injection, the antigen is slowly released from the injection site as alum particles slowly disaggregate over time (*27, 31, 42*). Using this simple approach, alum-adsorbed MD39 trimers that normally clear from the injection site within a few days instead clear much more slowly, over ∼10 days (*27*). We thus tested a two-dose ED immunization giving 20% of the vaccine dose on day 0 as a bolus and 80% of the dose as an alum/pSer-trimer formulation on day 7. Total GC B cells and Tfh cells elicited were similar between the 2-ED and 2-ED (alum-pSer) groups (**Fig. 6K-L**). However, relative to the bolus 2-ED immunization, extended antigen delivery provided by alum particle anchoring of the second shot in the 2-ED (alum-pSer) group trended towards improved responses, eliciting a 2.5-fold increase in the frequency of intact trimer-binding GC B cells and a 2-fold increase in the absolute number of these trimer-specific cells (**Fig. 6M-O**). These observations are consistent with model predictions (**Fig. 6E-F**). Interestingly, using the pSer-alum anchoring approach for both shots of the 2-dose regimen showed no improvement in trimer-specific GC responses over using it only for the 2^nd^ shot (**fig. S5A-C**) indicating that the benefit of constant release of antigen is more relevant once high antibody levels are present to enable capture of the native antigen. *In silico* simulation of this scenario agrees with this experimental observation (**fig. S5D-E**). Thus, an engineered 2-dose immunization, providing an initial “priming” dose followed by a larger extended-release vaccine dose retains much of the benefit to the GC response and fully amplifies the serum antibody response, substantially greater than bolus immunization.

## Discussion

Germinal center responses are critical to the evolution of diverse and high affinity antibody responses, and the size of the early GC response has been shown to predict the magnitude of neutralizing antibodies generated by HIV Env trimer immunization in rhesus macaques (*17*). In previous studies, we discovered that prolonging vaccine availability through extended dosing strategies such as implantable osmotic pumps or repeated injections of a given dose of vaccine over time alters multiple facets of the immune response– increasing the number of B cells entering GCs, the number of antigen-specific Tfh cells generated, and the number of unique clones in the GC, accompanied by increased total antibody titers, memory B cells, and neutralizing antibody production (*17*, *19*, *20*). In particular, an escalating-dose immunization of 7 shots administered in an exponentially-increasing dose pattern over two weeks was particularly effective in both mice and non-human primates for promoting humoral responses to HIV Env immunogens (*19*, *20*, *22*). However, administering seven doses is not practical for mass vaccination. Here we sought to better understand the critical elements of this potent dosing strategy, and based on our understanding of the mechanisms of ED-induced B cell responses, we hypothesized that a two-dose immunization, with an initial small dose of antigen to initiate B cell priming followed by a larger second dose that could be given to promote antigen deposition in follicles, might still offer substantial enhancements in humoral responses over traditional bolus vaccination. Such a scenario-two shots administered 1-2 weeks apart, would not be out of the realms of practicality (e.g., compare to current COVID-19 mRNA vaccines, administered twice at a 3- or 4-week interval).

Through systematic studies varying the number of doses, dose ratio, and dose intervals in mice, we found that a two-shot reduced ED regimen, administering 20% of the vaccine dose on day 0 and 80% on day 7 elicited 5-10-fold increases in peak antigen-binding total B cells (**Fig. 2B**) and GC B cells (**Fig. 1N**, **2E**), respectively, and increased serum antibody responses 60-fold relative to bolus immunization (**Fig. 2H**). Informed by computational modeling of the GC response, we further optimized an extended-release formulation on the 2^nd^ shot by anchoring the antigen onto alum via a phosphoserine linker resulting in even better antigen-specific GC responses over the reduced 2-shot regimen (**Fig. 6N-O**).

By combining computational modeling and experimental studies, we found that the motivating initial hypothesis was correct, wherein 2-ED priming promotes substantial antigen capture on FDCs, compared to very limited to undetectable antigen capture following bolus immunization. However, this is not the only effect of the extended-prime dosing. 2-ED immunization also increased Tfh responses (by 6-fold over bolus at the peak of the response, **Fig. 2D**). Tfh cells are essential for the GC reaction, providing support to B cells to generate long-lived plasma cells and memory B cells (*43–45*). The development of Tfh cells is initiated by priming of naïve T cells by DCs (*45*). The 2-shot immunization leads to a rapid increase in MHC-II^+^ DCs after the 2nd dose, which may continue the expansion of T cells initially primed after the first dose (*46*, *47*). The 2-shot regimen may also benefit from the fact that T cells begin to concentrate in GCs 5-8 days post immunization (*48*, *49*). Moreover, while we have abstracted this detail in our model, maintenance of the Tfh cell phenotype requires sustained antigenic stimulation (*33*), which is likely enhanced by increased antigen capture achieved in the 2-ED and 7-ED vaccinations.

Our synergistic deployment of computational and experimental studies has led to new insights into the fundamental dynamics of antibody development & antigen capture that can have broad implications designing vaccine dosing regimens. With extended dosing, availability of intact antigen in the lymph node is synchronized with the developing GC response. We observed that for the 7-ED regimen, amplification of GC and Tfh cells occurs towards the end of the dosing schedule, corresponding to the timing when antigen capture on FDCs is most prominent. The computational model suggests that an important mechanism underlying the efficacy of extended dosing regimens is the improved capture of native antigen on FDCs. In a simplified 2-shot immunization, administering a majority of the antigen on the 2^nd^ shot allows for exploitation of pre-existing antibody responses induced by the lower 1^st^ dose, maximizing the quantity of the immunogen dose captured in immune complexes. The computational model further predicts that if antigen is slowly released at 2^nd^ dose, increased representation of antigen in native form on FDC can further promote intact antigen-specific GC responses. This is in contrast to bolus immunization where only a small amount of antigen is presented on FDC to GC B cells, much of which is non-native protein and breakdown products that can be immunodominant and distract the GC response from relevant targets (*50*, *51*). These predictions were positively tested by our experiments. More antigen on FDCs during later stages of a GC likely increases GC B cell clonotypic diversity, allowing for significantly more BCR sequence space to be explored for high affinity BCR mutations.

Several technologies are being developed to recapitulate the effects of the 7-dose ED regimen. Our findings provide a strong rationale for utilizing reduced dosing immunization strategies that promote native antigen capture in the FDC network and enhance humoral responses. Developing a single-shot system that could mimic the effects of extended dosing would greatly improve compliance and increase vaccine coverage in developing countries. We have demonstrated the potential of a 2-shot regimen, but several single-shot approaches could also achieve this goal. For example, biodegradable polymer formulations have been demonstrated that achieve programmed timing of vaccine release at the injection site (*52*), and slow-dissolving alum coatings have been developed that may enable vaccine to be released over tunable time periods *in vivo* (*28*). These technologies could be particularly important for generating protective antibody responses against challenging pathogens such as HIV and for generating broadly neutralizing antibody responses against other variable pathogens. However, it is crucial to consider potential toxicities associated with these regimens, as multiple immunizations of potent adjuvants could result in undesirable side effects. Similarly, extended delivery technologies must avoid injection site reactogenicity and chronic inflammation. Collectively, these efforts will establish effective strategies that can be broadly applied in vaccine design to achieve the benefits of extended dosing in real-world vaccines. A limitation of this study is that immune response dynamics, and particularly the kinetics of the GC response, may differ in mice vs. non-human primates or humans. We expect the key elements of this strategy to be valid across species, but the ideal dosing ratio and dosing interval for a 2-ED regimen in NHPs or humans will require experimental evaluation.

In summary, we have identified optimized extended two-dose extended priming approaches that greatly improve multiple facets of the humoral immune responses to HIV immunogens compared to traditional bolus vaccination. Extended prime immunizations of this type augment GC and Tfh responses, increase trimer-specific serum antibody titers over bolus immunization; and highlight the critical role of antigen capture dynamics and priming of DCs in driving the enhanced immune responses.

## MATERIALS & METHODS

### Study design

The primary aim of this research was to assess the impact of extended prime immunizations on humoral responses to subunit vaccine immunization, compared to traditional bolus immunization in mouse models. To achieve this goal, we immunized mice with clinically relevant subunit protein immunogens in combination with saponin adjuvants, and evaluated early (antigen uptake and induction of Tfh and GC) and late (lymph node and serum antibody) responses over a period of time. The mechanistic studies conducted focused on examining antigen acquisition and activation of antigen-specific B and T cells, as well as computational analyses of the germinal center response to parse out differences in the dosing regimens.

### Immunogens

N332-GT2 trimers were expressed in FreeStyle 293F cells (Invitrogen, Cat no. R79007) and purified in two steps by affinity chromatography using a GE HisTrap column and size-exclusion chromatography using a GE S200 Increase column as described previously (*29*, *53*). MD39 HIV Env trimer was generated as previously described (*53*). Both trimers were administered at a dose of 10 ug per animal.

### pSer-conjugation

MD39 trimer immunogens bearing a free C-terminal cysteine at a concentration of 1 mg/ml were reduced by incubation with 10 molar equivalents of tris(2-carboxyethyl)phosphine (TCEP, ThermoFisher), followed by incubation at 25°C for 10 minutes. The reduced protein solutions were then processed using Amicon Ultra Centrifugal Filters (10 kDa MWCO, Millipore Sigma) to remove TCEP, and the resulting protein was mixed with 5 molar equivalents of pSer-maleimide linkers at a concentration of 1 mg/ml for 16 hours at 4°C in tris-buffered saline (TBS, Sigma Aldrich) at pH 7.2-7.4. After the reaction, unreacted pSer linker was removed using centrifugal filters in TBS, and pSer-antigen was buffer exchanged to PBS.

### Adjuvant preparation

The saponin adjuvant used in this study, SMNP, is an ISCOM-like self-assembled nanoparticle consisting of Quillaja saponin, cholesterol, DPPC, and MPLA (*30*). Briefly, solutions of cholesterol (20mg/ml final concentration, Avanti Polar Lipids Cat# 700000), DPPC (20mg/ml final concentration, Avanti Polar Lipids Cat# 850355), and MPLA (10mg/ml final concentration, PHAD) were prepared in Milli-Q water containing 20% w/vol MEGA-10 (Sigma D6277) detergent. Quil-A saponin (InvivoGen vac-quil) was dissolved in Milli-Q water at a final concentration of 100 mg/ml. All components were mixed at a molar ratio of 10:10:2.5:1 (Quil-A:chol:DPPC:MPLA) followed by dilution with PBS to a final concentration of 1 mg/ml cholesterol. The solution was allowed to equilibrate overnight at room temperature, followed by dialysis against PBS using a 10k MWCO membrane. The adjuvant solution was then sterile filtered, concentrated using 50k MWCO centricon spin filters, and further purified by FPLC using a Sephacryl S-500 HR size exclusion column. Doses are reported in terms of the amount of saponin administered, calculated by measuring the concentration of cholesterol (Cholesterol Quantitation kit; Millipore Sigma; Cat# MAK043) in the preparation and assuming quantitative incorporation of the saponin during synthesis.

### Antigen labeling and characterization

A 1 mg/mL solution of protein antigen (N332-GT2) in PBS was mixed with an equal volume of 0.2 M sodium bicarbonate buffer (pH 8.4) on ice. A fresh stock solution of Sulfo-Cyanine 5 NHS ester was prepared at a concentration of 1 mg/mL in 0.2 M sodium bicarbonate buffer (pH 8.4) and added to the antigen solution. The mixture was incubated at 4°C for 16 hours, and then desalted twice using a Zeba Spin Desalting column equilibrated in PBS. The labeled antigen was filtered through 0.22 µm pore size Spin-X centrifuge tube filters and stored at 4°C until use. The degree of labeling of the antigen was determined by measuring the absorbance at 280 and 646 nm wavelengths for total protein and Cy5 dye, respectively. Extinction coefficient values of 113215 and 271000 M^-1^cm^-1^ were used to calculate the relative concentrations of one subunit of N332-GT2 Trimer and sulfo-cy5 NHS ester, respectively. The degree of labeling for soluble antigen was determined by calculating the ratio of antigen concentration to Cy5 concentration.

### Mice

All mouse studies were performed according to institutional and National Institutes of Health (NIH) guidelines for humane animal use and in accordance with the Association for Assessment and Accreditation of Laboratory Animal Care. Protocols were approved by the Institutional Animal Care and Use Committee (IACUC) at MIT. All *in vivo* experiments were performed in 8-week-old female C57Bl/6J mice (Jackson Laboratory). Experiments were performed in specific pathogen-free animal facilities at the MIT Koch Institute for Integrative Cancer Research. Mice were housed under standard 12-hour light - 12-hour dark conditions with ad libitum access to water and chow.

### Immunizations and sample collections

8-week-old female C57BL/6 mice were anesthetized and immunized with 10 µg of indicated antigens (N332-GT2 trimer or MD39 trimer) in the presence of 5 µg saponin adjuvant (SMNP) subcutaneously, with half of the dose administered on each side of the tail base. In the case of the pSer-conjugated MD39 trimer antigen, immunizations were prepared by mixing 10 μg of antigen and 50 μg of alum in 100 μl of sterile TBS (Sigma-Aldrich Alhydrogel, catalog no. T5912) per mouse unless otherwise specified. Antigen was loaded onto alum for 30 min on a tube rotator after which 5 μg of SMNP was added into the immunization and incubated with antigen-alum formulations for 30 min before immunization. This dose of SMNP corresponds to 5 μg of Quil-A and 0.5 μg of MPLA. Blood (from submandibular; 100 μL) was collected at indicated time-points into serum separator tubes (BD Corporation) and centrifuged at 4,000 × g for 10 min at 4 °C. Sera extracted from blood samples were stored at −80 °C until ready for analysis. Inguinal LNs were harvested and added to Eppendorf tubes containing Protease inhibitor buffer (containing protease inhibitor cocktail and EDTA in PBS with 2% FBS). LNs were processed using a pestle and centrifuged at 16,000g for 5 min at 4 °C to pellet the cell/tissue debris. Supernatant was transferred to Spin-X tubes (Corning™ Costar™) and centrifuged again for 5 min with the flow through being transferred to final collection tubes, flash frozen and stored at −80 °C until ready for analysis.

### ELISA

To analyze on-target antibody response, high-binding ELISA plates (07-200-37, Fisher Scientific) were coated with 1 mg/ml trimer and blocked with 2% BSA in PBS overnight. To detect antigen-specific IgG responses, dilutions of serum or lymph node aspirate were added to the wells and incubated for 1.5 hours at 25°C. Plates were washed three times in PBS containing 0.2% Tween-20, and then anti-mouse IgG secondary antibody conjugated to HRP (172-1011, Bio-Rad Laboratories), diluted 1:5000 in blocking buffer as per manufacturer instructions, was added to the wells. After 1 hour of incubation, plates were again washed, developed with TMB, and the reaction was stopped with sulfuric acid. The optical density of the mixture was read out at 450 nm minus the absorbance at 540 nm according to the manufacturer’s instructions.

### Immunofluorescence staining

Inguinal lymph nodes (LNs) extracted from euthanized mice were submerged into cryomolds containing O.C.T. (23-730-571, Fisher Scientific) compound and dipped into 2-methylbutane (M32631, Millipore Sigma) pre-chilled in liquid nitrogen. All frozen tissues were cryosectioned on a Leica CM1950 at 10 µm thickness, adhered to Superfrost Plus microscope slides (12-550-15, Fisher Scientific), and stored in −80°C until use. Frozen sections were retrieved from −80°C, quickly thawed, and incubated in 4% paraformaldehyde for 10 minutes at 25°C. The sections were washed 3 times in PBS with 10-minute incubation time between each wash. Excess PBS was removed after the final wash before incubating the slides in blocking buffer, comprised of 2% BSA and 2% Triton X-100 in PBS. After 30 minutes, the blocking buffer was aspirated and the slides were stained in 1:75 anti-CD35 (740029, BD Biosciences) primary antibody solution also made in blocking buffer for ∼16 hours at 4°C. These slides were washed in PBS 3 times for 10 min, stained with 1:200 diluted secondary antibodies solution in blocking buffer (ab150063 Abcam, ab150061, Abcam) for 4 hours at room temperature, and washed again in PBS. To mount the slides, one drop of ProLong Diamond Antifade Mountant (Thermo) was added directly onto the stained tissues prior to gently placing a 20x20 mm coverslip on top of the droplet to sandwich the section. The coverslip was sealed using CoverGrip coverslip sealant (23005, Biotium) and imaged immediately. For all experiments, imaging was performed on a Leica SP8 confocal microscope equipped with a white light laser and spectral emission filter to detect emission wavelengths between 470 and 670 nm with a minimum bandwidth of 10 nm. All images were recorded with a 25X water immersion lens and a 63X oil immersion lens for assessing antigen drainage in the LNs, laser settings were kept constant across different time points for each immunogen.

### Whole tissue imaging

For whole tissue imaging, mice were injected with anti-CD35 BV421 antibody (clone 8C12) and lymph nodes were isolated after 16 hours and fixed overnight in 4% paraformaldehyde. Lymph nodes were then processed with a modified DISCO protocol as previously described (*54*). Briefly, the nodes were washed twice in PBS and excess fat and connective tissue was removed. Nodes were then gradually moved into solutions containing successively high concentrations (20, 50, 80%) of methanol for 30 mins, until they were incubated for half an hour in pure methanol. Nodes were then bleached for 2 minutes in hydrogen peroxide and returned to methanol for half an hour. They were then gradually moved into solutions containing increasing concentrations of tertiary-butanol (20, 50, 80%) before eventually being incubated in pure tertiary-butanol for one hour at 37° C. All solutions used after bleaching contained an additional 0.4% α-tocopherol (vitamin E). Nodes were then removed from solution and allowed to dry completely before being placed in dichloromethane for delipidation. After the 8 lymph nodes dropped to the bottom of tubes following swirling (indicating removal of remaining tertiary-butanol), they were stored in dibenzyl ether with 0.4% α-tocopherol, which was used as a refractive index matching solution. Cleared lymph nodes were imaged using an Olympus FV1200 Laser Scanning Confocal Microscope at 10x magnification. Lasers were set to minimize pixel saturation in the brightest samples. All laser and channel settings were then kept constant across groups for direct comparison between different samples. Each lymph node was imaged over a depth of 300 µm with line average of 3. FDC occupancy calculations were performed with a MATLAB script, where images for each channel were smoothed with a 3-D Gaussian filter (sigma = 0.5), then binarized into a mask to identify follicle or antigen area. The fraction of FDC area occupied by antigen was achieved by calculating overlapping pixels in the two binary masks. This calculation was performed for each individual image in the z-stack (9 per image), as well as for the z-projection (sum of slices).

### Flow Cytometry Analysis of Lymph Nodes

Inguinal lymph nodes were harvested, and single-cell suspensions were obtained by passage of the lymph nodes through a 70-μm filter (BD Biosciences). The isolated cells were stained with Live/Dead fixable aqua stain (L34957, Thermo Fisher Scientific) for 10 min at 25°C before washing twice in flow cytometry buffer. Cells were then incubated with Fc block for 10 min at 4°C before staining with antibodies listed (see the supplementary materials) for 20 additional min at 4°C. For trimer specific GC B cell analysis, cells stained with antibodies were distributed evenly and exposed to biotinylated trimer preincubated with streptavidin (30 min at molar ratio of 1:4 at 25°C) conjugated to phycoerythrin (405203, BioLegend) and/or BV421 (405226, BioLegend). Flow cytometry was carried out on a BD LSR Fortessa or LSR II. To assess GC B cells, cells were stained with GL7 PerCPCy5.5 (GL7; BioLegend), CD38 FITC (90; BioLegend), B220 PE-Cy7 (RA3-6B2; BioLegend), and CD4 BV711 (GK1.5; BioLegend). To assess Tfh cells, cells were stained with CD4 BV711 (GK1.5; BioLegend), B220 PE-Cy7 (RA3-6B2; BioLegend), CXCR5 PE (phycoerythrin) (L138D7; BioLegend), and PD1 BV421 (29F.1A12; BioLegend). Dead cells were stained using Fixable Aqua Dead Cell Stain kit (Thermo Fisher Scientific).

### Statistical analysis

Statistical analysis and graphing were done with GraphPad Prism for the experimental data and with MATLAB Statistics and Machine Learning Toolbox for the simulation data. The two-tailed Student’s *t* test was used to compare two experimental groups and one-way Anova with Dunnett’s multiple comparison post hoc analysis was used for comparing more than two groups. Details of the statistical test and number of replicates are indicated in the figure legends. A value of *P*<0.05 was considered statistically significant.

### Mathematical model

The coarse-grained model of T cell priming by antigen and adjuvant was formulated by integrating insights from experimental literature (*30*, *55–60*) with a previously published model of T cell proliferation (*37*). The system of ordinary differential equations is summarized in Fig. S2A, and a detailed description of the model and parameters is provided in Supplementary Materials. The model for the GC B cell response was adapted with minor modifications from a previous published study (*35*). A comprehensive overview of the model, including the equations, parameters, and the details of the modifications made, is provided in the Supplementary Materials. The MATLAB scripts and instructions to reproduce the results can be found on our GitHub repository at github.com/leerang77/Extended_Priming.

## Acknowledgements

We thank the Koch Institute’s Robert A. Swanson (1969) Biotechnology Center for technical support, specifically the Flow Cytometry and Microscopy Facilities. This work was supported in part by the Koch Institute Core Grant (P30-CA14051) from the National Cancer Institute, the NIH (awards AI161297, AI125068, and UM1AI144462 to DJI and AI175489 to DJI and AKC), and the Ragon Institute of MIT, MGH, and Harvard. Leerang Yang was also supported by NIH grant # U19AI057229

## Author contributions

S.H.B, D.J.I, A.K,C and L.Y conceived the experiments and computational modeling framework, wrote the manuscript together and all authors discussed and commented on it. S.H.B, L.M, E.B.A, K.A.R, A.M.R, H.S, A.A, S.W, and A.W. performed all the experimental work and analyzed the data. L.Y. performed all the computational work, and interpreted them with A.K.C, S.H.B. and D.J.I.. S.H.B and L.Y generated the figures. D.J.I and A.K.C acquired funding and supervised the research.

## REFERENCES

1. D. H. Barouch, Covid-19 Vaccines — Immunity, Variants, Boosters. New Engl J Med. 387, 1011– 1020 (2022).

2. J. H. Lee, S. Crotty, HIV vaccinology: 2021 update. Semin Immunol. 51, 101470 (2021).

3. B. F. Haynes, K. Wiehe, P. Borrow, K. O. Saunders, B. Korber, K. Wagh, A. J. McMichael, G. Kelsoe, B. H. Hahn, F. Alt, G. M. Shaw, Strategies for HIV-1 vaccines that induce broadly neutralizing antibodies. Nat Rev Immunol. 23, 142–158 (2023).

4. M. B. Feinberg, Uhambo — Twists and Turns on the Journey to an Efficacious HIV-1 Vaccine. New Engl J Med. 384, 1157–1159 (2021).

5. D. R. Burton, L. Hangartner, Broadly Neutralizing Antibodies to HIV and Their Role in Vaccine Design. Annu. Rev. Immunol. 34, 635–659 (2016).

6. D. R. Burton, Advancing an HIV vaccine; advancing vaccinology. Nat Rev Immunol. 19, 77–78 (2019).

7. P. B. Gilbert, Y. Huang, A. C. deCamp, S. Karuna, Y. Zhang, C. A. Magaret, E. E. Giorgi, B. Korber, P. T. Edlefsen, R. Rossenkhan, M. Juraska, E. Rudnicki, N. Kochar, Y. Huang, L. N. Carpp, D. H. Barouch, N. N. Mkhize, T. Hermanus, P. Kgagudi, V. Bekker, H. Kaldine, R. E. Mapengo, A. Eaton, E. Domin, C. West, W. Feng, H. Tang, K. E. Seaton, J. Heptinstall, C. Brackett, K. Chiong, G. D. Tomaras, P. Andrew, B. T. Mayer, D. B. Reeves, M. E. Sobieszczyk, N. Garrett, J. Sanchez, C. Gay, J. Makhema, C. Williamson, J. I. Mullins, J. Hural, M. S. Cohen, L. Corey, D. C. Montefiori, L. Morris, Neutralization titer biomarker for antibody-mediated prevention of HIV-1 acquisition. Nat Med. 28, 1924–1932 (2022).

8. L. Corey, P. B. Gilbert, M. Juraska, D. C. Montefiori, L. Morris, S. T. Karuna, S. Edupuganti, N. M. Mgodi, A. C. deCamp, E. Rudnicki, Y. Huang, P. Gonzales, R. Cabello, C. Orrell, J. R. Lama, F. Laher, E. M. Lazarus, J. Sanchez, I. Frank, J. Hinojosa, M. E. Sobieszczyk, K. E. Marshall, P. G. Mukwekwerere, J. Makhema, L. R. Baden, J. I. Mullins, C. Williamson, J. Hural, M. J. McElrath, C. Bentley, S. Takuva, M. M. G. Lorenzo, D. N. Burns, N. Espy, A. K. Randhawa, N. Kochar, E. Piwowar-Manning, D. J. Donnell, N. Sista, P. Andrew, J. G. Kublin, G. Gray, J. E. Ledgerwood, J. R. Mascola, M. S. Cohen, H. 704/HPTN 085 and H. 703/HPTN 081 S. Teams, Two Randomized Trials of Neutralizing Antibodies to Prevent HIV-1 Acquisition. New Engl J Med. 384, 1003–1014 (2021).

9. C. Havenar-Daughton, J. H. Lee, S. Crotty, Tfh cells and HIV bnAbs, an immunodominance model of the HIV neutralizing antibody generation problem. Immunol Rev. 275, 49–61 (2017).

10. P. J. Klasse, G. Ozorowski, R. W. Sanders, J. P. Moore, Env Exceptionalism: Why Are HIV-1 Env Glycoproteins Atypical Immunogens? Cell Host Microbe. 27, 507–518 (2020).

11. F. Krammer, P. Palese, Advances in the development of influenza virus vaccines. Nat. Rev. Drug Discov. 14, 167–182 (2015).

12. F. Krammer, P. Palese, Universal Influenza Virus Vaccines That Target the Conserved Hemagglutinin Stalk and Conserved Sites in the Head Domain. J. Infect. Dis. 219, S62–S67 (2019).

13. G. D. Victora, M. C. Nussenzweig, Germinal Centers. Annu Rev Immunol. 40, 1–30 (2022).

14. G. D. Victora, M. C. Nussenzweig, Germinal Centers. Immunology. 30, 429–457 (2012).

15. Z. Shulman, A. D. Gitlin, J. S. Weinstein, B. Lainez, E. Esplugues, R. A. Flavell, J. E. Craft, M. C. Nussenzweig, Dynamic signaling by T follicular helper cells during germinal center B cell selection. Science. 345, 1058–1062 (2014).

16. A. D. Gitlin, C. T. Mayer, T. Y. Oliveira, Z. Shulman, M. J. K. Jones, A. Koren, M. C. Nussenzweig, T cell help controls the speed of the cell cycle in germinal center B cells. Science. 349, 643–646 (2015).

17. M. Pauthner, C. Havenar-Daughton, D. Sok, J. P. Nkolola, R. Bastidas, A. V. Boopathy, D. G. Carnathan, A. Chandrashekar, K. M. Cirelli, C. A. Cottrell, A. M. Eroshkin, J. Guenaga, K. Kaushik, D. W. Kulp, J. Liu, L. E. McCoy, A. L. Oom, G. Ozorowski, K. W. Post, S. K. Sharma, J. M. Steichen, S. W. de Taeye, T. Tokatlian, A. T. de la Peña, S. T. Butera, C. C. LaBranche, D. C. Montefiori, G. Silvestri, I. A. Wilson, D. J. Irvine, R. W. Sanders, W. R. Schief, A. B. Ward, R. T. Wyatt, D. H. Barouch, S. Crotty, D. R. Burton, Elicitation of Robust Tier 2 Neutralizing Antibody Responses in Nonhuman Primates by HIV Envelope Trimer Immunization Using Optimized Approaches. Immunity. 46, 1073–1088.e6 (2017).

18. R. K. Abbott, J. H. Lee, S. Menis, P. Skog, M. Rossi, T. Ota, D. W. Kulp, D. Bhullar, O. Kalyuzhniy, C. Havenar-Daughton, W. R. Schief, D. Nemazee, S. Crotty, Precursor Frequency and Affinity Determine B Cell Competitive Fitness in Germinal Centers, Tested with Germline-Targeting HIV Vaccine Immunogens. Immunity. 48, 133–146.e6 (2018).

19. H. H. Tam, M. B. Melo, M. Kang, J. M. Pelet, V. M. Ruda, M. H. Foley, J. K. Hu, S. Kumari, J. Crampton, A. D. Baldeon, R. W. Sanders, J. P. Moore, S. Crotty, R. Langer, D. G. Anderson, A. K. Chakraborty, D. J. Irvine, Sustained antigen availability during germinal center initiation enhances antibody responses to vaccination. Proc National Acad Sci. 113, E6639–E6648 (2016).

20. K. M. Cirelli, D. G. Carnathan, B. Nogal, J. T. Martin, O. L. Rodriguez, A. A. Upadhyay, C. A. Enemuo, E. H. Gebru, Y. Choe, F. Viviano, C. Nakao, M. G. Pauthner, S. Reiss, C. A. Cottrell, M. L. Smith, R. Bastidas, W. Gibson, A. N. Wolabaugh, M. B. Melo, B. Cossette, V. Kumar, N. B. Patel, T. Tokatlian, S. Menis, D. W. Kulp, D. R. Burton, B. Murrell, W. R. Schief, S. E. Bosinger, A. B. Ward, C. T. Watson, G. Silvestri, D. J. Irvine, S. Crotty, Slow Delivery Immunization Enhances HIV Neutralizing Antibody and Germinal Center Responses via Modulation of Immunodominance. Cell. 177, 1153–1171.e28 (2019).

21. D. J. Irvine, A. Aung, M. Silva, Controlling timing and location in vaccines. Adv Drug Deliver Rev (2020), doi:10.1016/j.addr.2020.06.019.

22. J. H. Lee, H. J. Sutton, C. A. Cottrell, I. Phung, G. Ozorowski, L. M. Sewall, R. Nedellec, C. Nakao, M. Silva, S. T. Richey, J. L. Torres, W.-H. Lee, E. Georgeson, M. Kubitz, S. Hodges, T.-M. Mullen, Y. Adachi, K. M. Cirelli, A. Kaur, C. Allers, M. Fahlberg, B. F. Grasperge, J. P. Dufour, F. Schiro, P. P. Aye, O. Kalyuzhniy, A. Liguori, D. G. Carnathan, G. Silvestri, X. Shen, D. C. Montefiori, R. S. Veazey, A. B. Ward, L. Hangartner, D. R. Burton, D. J. Irvine, W. R. Schief, S. Crotty, Long-primed germinal centres with enduring affinity maturation and clonal migration. Nature. 609, 998–1004 (2022).

23. A. V. Boopathy, A. Mandal, D. W. Kulp, S. Menis, N. R. Bennett, H. C. Watkins, W. Wang, J. T. Martin, N. T. Thai, Y. He, W. R. Schief, P. T. Hammond, D. J. Irvine, Enhancing humoral immunity via sustained-release implantable microneedle patch vaccination. Proc National Acad Sci. 116, 16473– 16478 (2019).

24. G. A. Roth, E. C. Gale, M. Alcántara-Hernández, W. Luo, E. Axpe, R. Verma, Q. Yin, A. C. Yu, H. L. Hernandez, C. L. Maikawa, A. A. A. Smith, M. M. Davis, B. Pulendran, J. Idoyaga, E. A. Appel, Injectable Hydrogels for Sustained Codelivery of Subunit Vaccines Enhance Humoral Immunity. Acs Central Sci. 6, 1800–1812 (2020).

25. E. C. Gale, A. E. Powell, G. A. Roth, E. L. Meany, J. Yan, B. S. Ou, A. K. Grosskopf, J. Adamska, V. C. T. M. Picece, A. I. d’Aquino, B. Pulendran, P. S. Kim, E. A. Appel, Hydrogel-Based Slow Release of a Receptor Binding Domain Subunit Vaccine Elicits Neutralizing Antibody Responses Against SARS-CoV-2. Adv. Mater. 33, 2104362 (2021).

26. G. A. Roth, V. C. T. M. Picece, B. S. Ou, W. Luo, B. Pulendran, E. A. Appel, Designing spatial and temporal control of vaccine responses. Nat Rev Mater, 1–22 (2021).

27. T. J. Moyer, Y. Kato, W. Abraham, J. Y. H. Chang, D. W. Kulp, N. Watson, H. L. Turner, S. Menis, R. K. Abbott, J. N. Bhiman, M. B. Melo, H. A. Simon, S. H.-D. la Mata, S. Liang, G. Seumois, Y. Agarwal, N. Li, D. R. Burton, A. B. Ward, W. R. Schief, S. Crotty, D. J. Irvine, Engineered immunogen binding to alum adjuvant enhances humoral immunity. Nat Med. 26, 430–440 (2020).

28. R. L. Garcea, N. M. Meinerz, M. Dong, H. Funke, S. Ghazvini, T. W. Randolph, Single-administration, thermostable human papillomavirus vaccines prepared with atomic layer deposition technology. Npj Vaccines. 5, 45 (2020).

29. J. M. Steichen, Y.-C. Lin, C. Havenar-Daughton, S. Pecetta, G. Ozorowski, J. R. Willis, L. Toy, D. Sok, A. Liguori, S. Kratochvil, J. L. Torres, O. Kalyuzhniy, E. Melzi, D. W. Kulp, S. Raemisch, X. Hu, S. M. Bernard, E. Georgeson, N. Phelps, Y. Adachi, M. Kubitz, E. Landais, J. Umotoy, A. Robinson, B. Briney, I. A. Wilson, D. R. Burton, A. B. Ward, S. Crotty, F. D. Batista, W. R. Schief, A generalized HIV vaccine design strategy for priming of broadly neutralizing antibody responses. Science. 366, eaax4380 (2019).

30. M. Silva, Y. Kato, M. B. Melo, I. Phung, B. L. Freeman, Z. Li, K. Roh, J. W. V. Wijnbergen, H. Watkins, C. A. Enemuo, B. L. Hartwell, J. Y. H. Chang, S. Xiao, K. A. Rodrigues, K. M. Cirelli, N. Li, S. Haupt, A. Aung, B. Cossette, W. Abraham, S. Kataria, R. Bastidas, J. Bhiman, C. Linde, N. I. Bloom, B. Groschel, E. Georgeson, N. Phelps, A. Thomas, J. Bals, D. G. Carnathan, D. Lingwood, D. R. Burton, G. Alter, T. P. Padera, A. M. Belcher, W. R. Schief, G. Silvestri, R. M. Ruprecht, S. Crotty, D. J. Irvine, A particulate saponin/TLR agonist vaccine adjuvant alters lymph flow and modulates adaptive immunity. Sci Immunol. 6, eabf1152 (2021).

31. K. A. Rodrigues, S. A. Rodriguez-Aponte, N. C. Dalvie, J. H. Lee, W. Abraham, D. G. Carnathan, L. E. Jimenez, J. T. Ngo, J. Y. H. Chang, Z. Zhang, J. Yu, A. Chang, C. Nakao, B. Goodwin, C. A. Naranjo, L. Zhang, M. Silva, D. H. Barouch, G. Silvestri, S. Crotty, J. C. Love, D. J. Irvine, Phosphate-mediated coanchoring of RBD immunogens and molecular adjuvants to alum potentiates humoral immunity against SARS-CoV-2. Sci Adv. 7, eabj6538 (2021).

32. C. Havenar-Daughton, D. G. Carnathan, A. T. de la Peña, M. Pauthner, B. Briney, S. M. Reiss, J. S. Wood, K. Kaushik, M. J. van Gils, S. L. Rosales, P. van der Woude, M. Locci, K. M. Le, S. W. de Taeye, D. Sok, A. U. R. Mohammed, J. Huang, S. Gumber, A. Garcia, S. P. Kasturi, B. Pulendran, J. P. Moore, R. Ahmed, G. Seumois, D. R. Burton, R. W. Sanders, G. Silvestri, S. Crotty, Direct Probing of Germinal Center Responses Reveals Immunological Features and Bottlenecks for Neutralizing Antibody Responses to HIV Env Trimer. Cell Reports. 17, 2195–2209 (2016).

33. D. Baumjohann, S. Preite, A. Reboldi, F. Ronchi, K. M. Ansel, A. Lanzavecchia, F. Sallusto, Persistent Antigen and Germinal Center B Cells Sustain T Follicular Helper Cell Responses and Phenotype. Immunity. 38, 596–605 (2013).

34. M. A. Mintz, J. G. Cyster, T follicular helper cells in germinal center B cell selection and lymphomagenesis. Immunol Rev. 296, 48–61 (2020).

35. L. Yang, M. V. Beek, Z. Wang, F. Muecksch, M. Canis, T. Hatziioannou, P. D. Bieniasz, M. C. Nussenzweig, A. K. Chakraborty, Antigen presentation dynamics shape the antibody response to variants like SARS-CoV-2 Omicron after multiple vaccinations with the original strain. Cell Reports. 42, 112256 (2022).

36. B. Pulendran, P. S. Arunachalam, D. T. O’Hagan, Emerging concepts in the science of vaccine adjuvants. Nat Rev Drug Discov, 1–22 (2021).

37. A. Mayer, Y. Zhang, A. S. Perelson, N. S. Wingreen, Regulation of T cell expansion by antigen presentation dynamics. Proc National Acad Sci. 116, 5914–5919 (2019).

38. P. Johansen, T. Storni, L. Rettig, Z. Qiu, A. Der-Sarkissian, K. A. Smith, V. Manolova, K. S. Lang, G. Senti, B. Müllhaupt, T. Gerlach, R. F. Speck, A. Bot, T. M. Kündig, Antigen kinetics determines immune reactivity. Proc National Acad Sci. 105, 5189–5194 (2008).

39. J. Quiel, S. Caucheteux, A. Laurence, N. J. Singh, G. Bocharov, S. Z. Ben-Sasson, Z. Grossman, W. E. Paul, Antigen-stimulated CD4 T-cell expansion is inversely and log-linearly related to precursor number. Proc National Acad Sci. 108, 3312–3317 (2011).

40. A. Aung, A. Cui, L. Maiorino, A. P. Amini, J. R. Gregory, M. Bukenya, Y. Zhang, H. Lee, C. A. Cottrell, D. M. Morgan, M. Silva, H. Suh, J. D. Kirkpatrick, P. Amlashi, T. Remba, L. M. Froehle, S. Xiao, W. Abraham, J. Adams, J. C. Love, P. Huyett, D. S. Kwon, N. Hacohen, W. R. Schief, S. N. Bhatia, D. J. Irvine, Low protease activity in B cell follicles promotes retention of intact antigens after immunization. Science. 379, eabn8934 (2023).

41. D. W. Kulp, J. M. Steichen, M. Pauthner, X. Hu, T. Schiffner, A. Liguori, C. A. Cottrell, C. Havenar-Daughton, G. Ozorowski, E. Georgeson, O. Kalyuzhniy, J. R. Willis, M. Kubitz, Y. Adachi, S. M. Reiss, M. Shin, N. de Val, A. B. Ward, S. Crotty, D. R. Burton, W. R. Schief, Structure-based design of native-like HIV-1 envelope trimers to silence non-neutralizing epitopes and eliminate CD4 binding. Nat Commun. 8, 1655 (2017).

42. K. A. Rodrigues, C. A. Cottrell, J. M. Steichen, B. Groschel, W. Abraham, H. Suh, Y. Agarwal, K. Ni, J. Y. H. Chang, P. Yousefpour, M. B. Melo, W. R. Schief, D. J. Irvine, Optimization of an alum-anchored clinical HIV vaccine candidate. npj Vaccines. 8, 117 (2023).

43. H. Qi, T follicular helper cells in space-time. Nat Rev Immunol. 16, 612–625 (2016).

44. S. Crotty, T Follicular Helper Cell Differentiation, Function, and Roles in Disease. Immunity. 41, 529–542 (2014).

45. S. Crotty, T Follicular Helper Cell Biology: A Decade of Discovery and Diseases. Immunity. 50, 1132–1148 (2019).

46. P. Devarajan, A. M. Vong, C. H. Castonguay, O. Kugler-Umana, B. L. Bautista, M. C. Jones, K. A. Kelly, J. Xia, S. L. Swain, Strong influenza-induced TFH generation requires CD4 effectors to recognize antigen locally and receive signals from continuing infection. Proc National Acad Sci. 119, e2111064119 (2022).

47. E. K. Deenick, A. Chan, C. S. Ma, D. Gatto, P. L. Schwartzberg, R. Brink, S. G. Tangye, Follicular Helper T Cell Differentiation Requires Continuous Antigen Presentation that Is Independent of Unique B Cell Signaling. Immunity. 33, 241–253 (2010).

48. D. Dominguez-Sola, G. D. Victora, C. Y. Ying, R. T. Phan, M. Saito, M. C. Nussenzweig, R. Dalla-Favera, The proto-oncogene MYC is required for selection in the germinal center and cyclic reentry. Nat Immunol. 13, 1083–1091 (2012).

49. Z. Shulman, A. D. Gitlin, S. Targ, M. Jankovic, G. Pasqual, M. C. Nussenzweig, G. D. Victora, T Follicular Helper Cell Dynamics in Germinal Centers. Science. 341, 673–677 (2013).

50. K. M. Cirelli, S. Crotty, Germinal center enhancement by extended antigen availability. Curr Opin Immunol. 47, 64–69 (2017).

51. J. K. Hu, J. C. Crampton, A. Cupo, T. Ketas, M. J. van Gils, K. Sliepen, S. W. de Taeye, D. Sok, G. Ozorowski, I. Deresa, R. Stanfield, A. B. Ward, D. R. Burton, P. J. Klasse, R. W. Sanders, J. P. Moore, S. Crotty, Murine Antibody Responses to Cleaved Soluble HIV-1 Envelope Trimers Are Highly Restricted in Specificity. J Virol. 89, 10383–10398 (2015).

52. A. K. J. McHugh, T. D. Nguyen, A. R. Linehan, D. Yang, A. M. Behrens, S. Rose, Z. L. Tochka, S. Y. Tzeng, J. J. Norman, A. C. Anselmo, X. Xu, S. Tomasic, M. A. Taylor, J. Lu, R. Guarecuco, R. Langer, A. Jaklenec, Fabrication of fillable microparticles and other complex 3D microstructures. Science. 357, 1138–1142 (2017).

52. J. M. Steichen, D. W. Kulp, T. Tokatlian, A. Escolano, P. Dosenovic, R. L. Stanfield, L. E. McCoy, G. Ozorowski, X. Hu, O. Kalyuzhniy, B. Briney, T. Schiffner, F. Garces, N. T. Freund, A. D. Gitlin, S. Menis, E. Georgeson, M. Kubitz, Y. Adachi, M. Jones, A. A. Mutafyan, D. S. Yun, C. T. Mayer, A. B. Ward, D. R. Burton, I. A. Wilson, D. J. Irvine, M. C. Nussenzweig, W. R. Schief, HIV Vaccine Design to Target Germline Precursors of Glycan-Dependent Broadly Neutralizing Antibodies. Immunity. 45, 483– 496 (2016).

53. T. Tokatlian, B. J. Read, C. A. Jones, D. W. Kulp, S. Menis, J. Y. H. Chang, J. M. Steichen, S. Kumari, J. D. Allen, E. L. Dane, A. Liguori, M. Sangesland, D. Lingwood, M. Crispin, W. R. Schief, D. J. Irvine, Innate immune recognition of glycans targets HIV nanoparticle immunogens to germinal centers. Science. 363, 649–654 (2019).

54. A. M. Didierlaurent, S. Morel, L. Lockman, S. L. Giannini, M. Bisteau, H. Carlsen, A. Kielland, O. Vosters, N. Vanderheyde, F. Schiavetti, D. Larocque, M. V. Mechelen, N. Garçon, AS04, an aluminum salt- and TLR4 agonist-based adjuvant system, induces a transient localized innate immune response leading to enhanced adaptive immunity. J. Immunol. (Baltim., MdJ: 1950). 183, 6186–97 (2009).

55. M. Dupuis, D. M. McDonald, G. Ott, Distribution of adjuvant MF59 and antigen gD2 after intramuscular injection in mice. Vaccine. 18, 434–439 (1999).

56. K. Murphy, C. Weaver, Janeway’s Immunobiology (2016), doi:10.1201/9781315533247.

57. S. Awate, L. A. Babiuk, G. Mutwiri, Mechanisms of Action of Adjuvants. Front. Immunol. 4, 114 (2013).

58. F. Liang, G. Lindgren, K. J. Sandgren, E. A. Thompson, J. R. Francica, A. Seubert, E. D. Gregorio, S. Barnett, D. T. O’Hagan, N. J. Sullivan, R. A. Koup, R. A. Seder, K. Loré, Vaccine priming is restricted to draining lymph nodes and controlled by adjuvant-mediated antigen uptake. Sci. Transl. Med. 9 (2017), doi:10.1126/scitranslmed.aal2094.

59. S. Calabro, M. Tortoli, B. C. Baudner, A. Pacitto, M. Cortese, D. T. O’Hagan, E. D. Gregorio, A. Seubert, A. Wack, Vaccine adjuvants alum and MF59 induce rapid recruitment of neutrophils and monocytes that participate in antigen transport to draining lymph nodes. Vaccine. 29, 1812–1823 (2011).

60. G. Altan-Bonnet, T. Mora, A. M. Walczak, Quantitative immunology for physicists. Phys. Rep. 849, 1–83 (2020).

61. T. Okada, M. J. Miller, I. Parker, M. F. Krummel, M. Neighbors, S. B. Hartley, A. O’Garra, M. D. Cahalan, J. G. Cyster, Antigen-Engaged B Cells Undergo Chemotaxis toward the T Zone and Form Motile Conjugates with Helper T Cells. PLoS Biol. 3, e150 (2005).

62. J. J. Taylor, K. A. Pape, H. R. Steach, M. K. Jenkins, Apoptosis and antigen affinity limit effector cell differentiation of a single naïve B cell. Science. 347, 784–787 (2015).

63. D. M.-Y. Sze, K.-M. Toellner, C. G. de Vinuesa, D. R. Taylor, I. C. M. MacLennan, Intrinsic Constraint on Plasmablast Growth and Extrinsic Limits of Plasma Cell Survival. J. Exp. Med. 192, 813–822 (2000).

64. I. Moran, A. Nguyen, W. H. Khoo, D. Butt, K. Bourne, C. Young, J. R. Hermes, M. Biro, G. Gracie, C. S. Ma, C. M. L. Munier, F. Luciani, J. Zaunders, A. Parker, A. D. Kelleher, S. G. Tangye, P. I. Croucher, R. Brink, M. N. Read, T. G. Phan, Memory B cells are reactivated in subcapsular proliferative foci of lymph nodes. Nat. Commun. 9, 3372 (2018).

65. M. V. Beek, M. C. Nussenzweig, A. K. Chakraborty, Two complementary features of humoral immune memory confer protection against the same or variant antigens. Proc. Natl. Acad. Sci. 119, e2205598119 (2022).

