## Supplementary Material for "Two-dose “extended priming” immunization amplifies humoral immune responses by synchronizing vaccine delivery with the germinal center response"

**The PDF file includes:**

**Figures S1 to S5**

**Table S1 to S3**

**Details of computational models**

**Other Supplementary Material for this manuscript includes the following:**

**Data file S1**

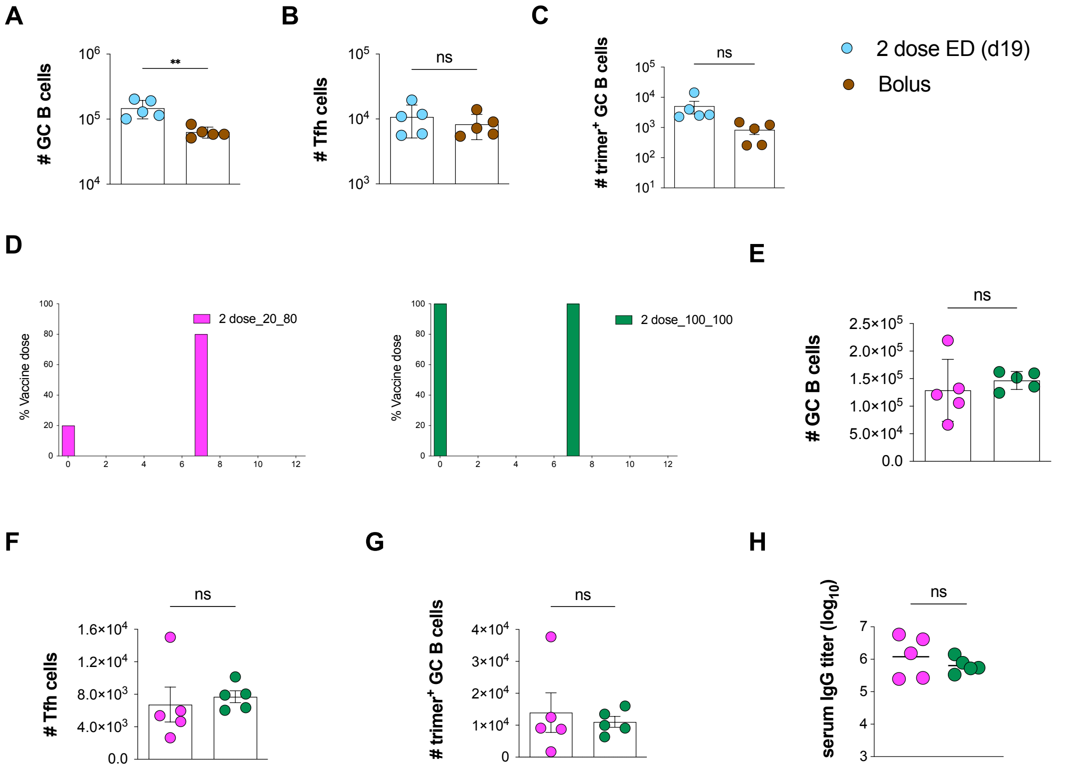


**figure S1: Additional 2-shot prime immunization GC response comparisons. (A-C)** C57BL/6J mice ($n$=5 animals/group) were immunized with 10 μg N332-GT2 trimer and 5 μg SMNP adjuvant either as a bolus immunization or 2-dose ED regimen (d0, d12) outlined in Fig 1A. Shown are flow cytometry analyses enumerating GC B cells **(A)**, Tfh cells **(B)**, and trimer-specific GC B cell responses **(C)** at day 19 for the 2-dose regimen compared to bolus immunization. **(D)** Schematic of 2-shot regimens comparing dose 1 (20%) and dose 2 (80%) vs dose 1 (100%) and dose 2 (100%) at day 14. **(E-H)** C57BL/6J mice ($n$=5 animals/group) were immunized with 10 μg N332-GT2 trimer and 5 μg SMNP adjuvant according to the dosing schemes in **(D)**. Shown are number of GC B cells **(E)**, Tfh cells **(F)**, and trimer-specific GC B cells **(G)** at day 14 and trimer-specific IgG titers **(H)** at day 28 for the two groups. Shown are data from one independent experiment for each immunization series. ****, $p < 0.0001$; ***, $p < 0.001$; **, $p < 0.01$; *, $p < 0.05$; ns, not significant; by one-way ANOVA with Dunnett's multiple comparison post test compared to bolus immunization.

**A**

$$\frac{d[Adj]}{dt} = -d_{Adj}[Adj]$$

$$\frac{d[TC]}{dt} = \frac{[Adj]}{S_{Adj} + [Adj]} - \mu[TC]$$

$$\frac{d[DC]}{dt} = D_0[TC] - \left(1 + k\frac{[Adj]}{S_{Adj} + [Adj]}\right)[DC][Ag] - \mu[DC]$$

$$\frac{d[Ag]}{dt} = -d_{Ag}[Ag]$$

$$\frac{d[aDC^{Ag^+}]}{dt} = \left(1 + k\frac{[Adj]}{S_{Adj} + [Adj]}\right)[DC][Ag] - \mu[aDC^{Ag^+}]$$

$$\frac{d[T]}{dt} = \alpha\frac{[aDC^{Ag^+}][T]}{[T] + [aDC^{Ag^+}]} - \eta([T] - T_0)$$

$$\frac{d[T_{FH}]}{dt} = \eta([T] - T_0)$$

| | |
|---|---|
| $d_{Ag}$ | Antigen decay rate |
| $d_{Adj}$ | Adjuvant decay rate |
| $S_{Adj}$ | Half-max adjuvant activity concentration |
| $k$ | Max increase in antigen uptake rate |
| $\alpha$ | Max T cell proliferation rate |
| $\eta$ | T cell differentiation rate |
| $\mu$ | Innate immune cell death rate |
| $D_0$ | Rate of DC recruitment |
| $T_0$ | Baseline number of T cells |

$Adj$ – Adjuvant, $TC$ – Tissue cells, $DC$ – Dendritic cells, $Ag$ – Antigen
$DC^{Ag^+}$ - Antigen-primed dendritic cells, $T$ – Proliferating T cells, $T_{FH}$ - Follicular helper T cells

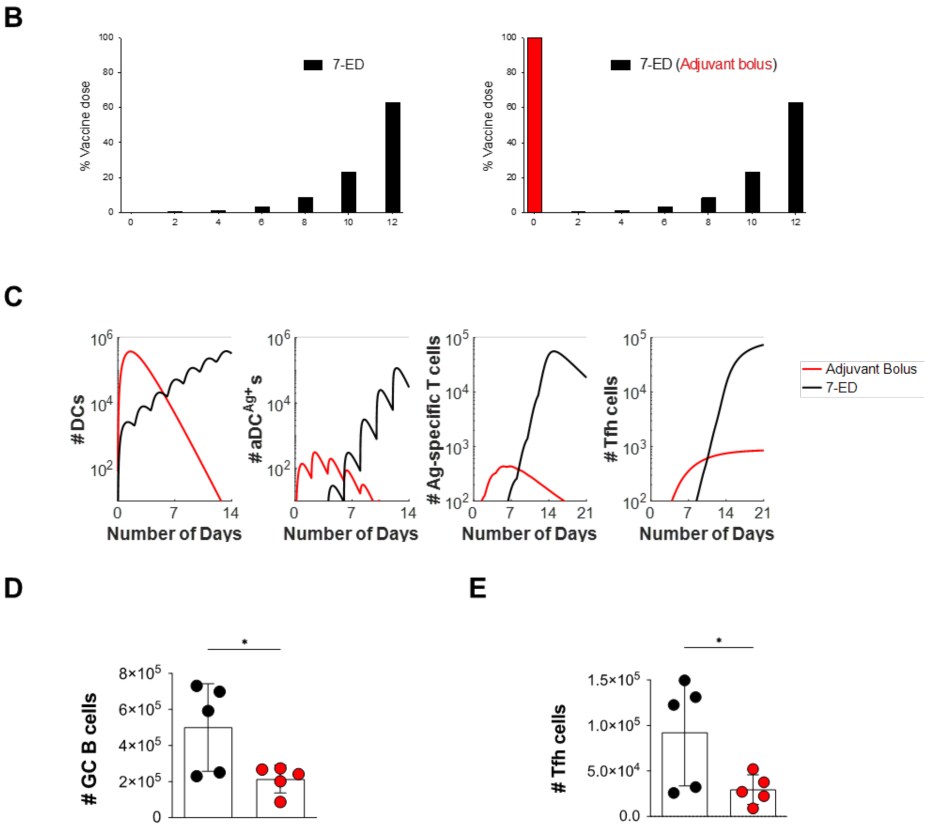


**figure S2: Computational model of T cell priming and analysis of a 7-ED (bolus adjuvant) regimen.** **(A)** Equations governing the kinetic model for innate immune responses and T cell priming. **(B)** Dosing scheme for 7-ED and 7-ED (adjuvant bolus) immunization regimens. **(C)** Model predictions for DC, $aDC^{Ag+}$, T cells, and Tfh cell responses under the standard 7-ED and 7-ED (adjuvant bolus) immunization schemes. **(D-E)** C57BL/6J mice ($n$=5 animals/group) were immunized with 10 µg N332-GT2 trimer and 5 µg SMNP adjuvant according to the dosing schemes in **(B)**. Shown is flow cytometry analysis on day 14 enumerating GC B cells **(D)** and Tfh cells **(E)**. Data from one independent experiment for each immunization series. ****, p < 0.0001; ***, p < 0.001; **, p < 0.01; *, p < 0.05; ns, not significant; by one-way ANOVA with Dunnett's multiple comparison post test compared to bolus immunization.

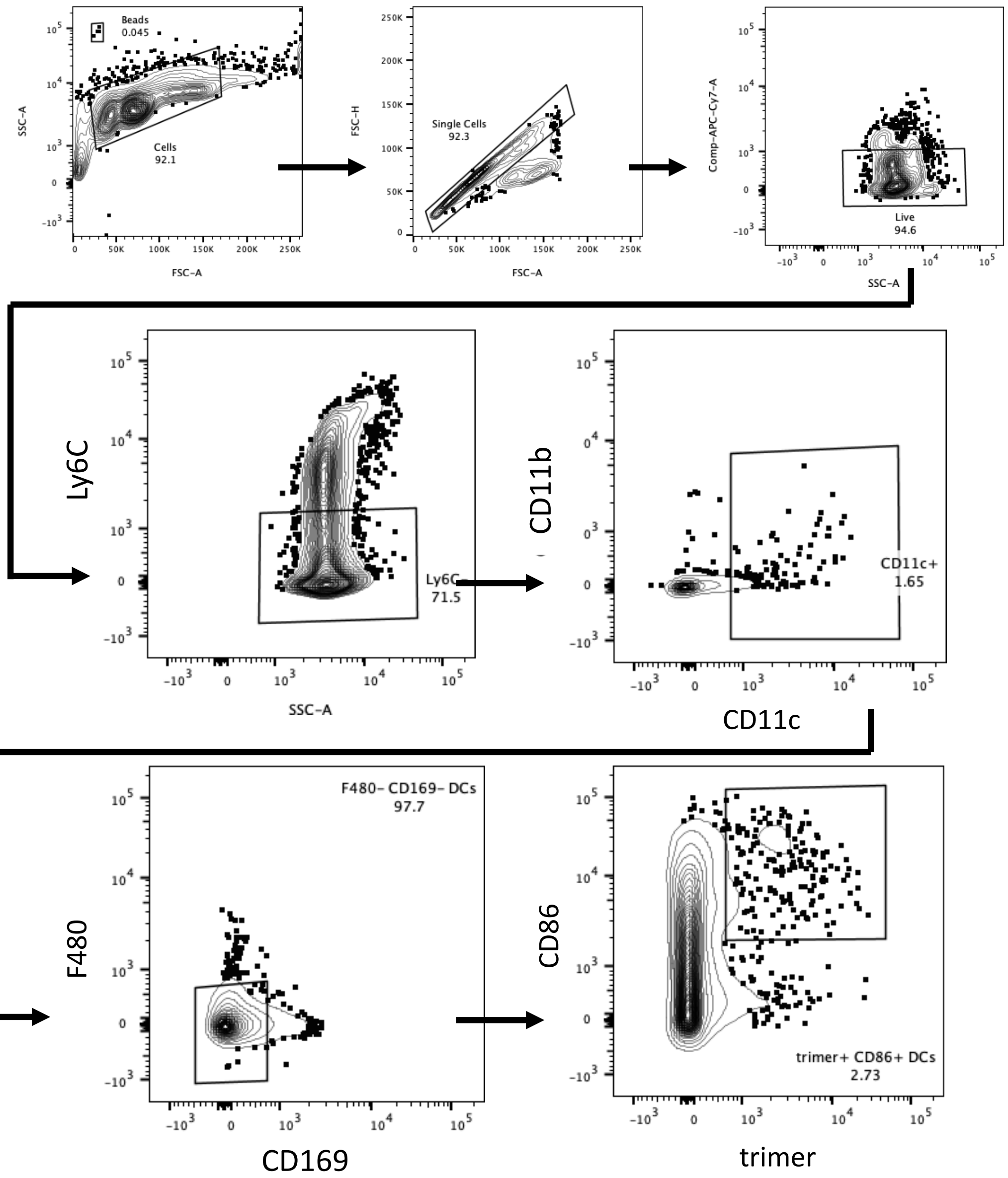


**figure S3: Gating strategy for DC kinetic analysis of different immunization regimens.**

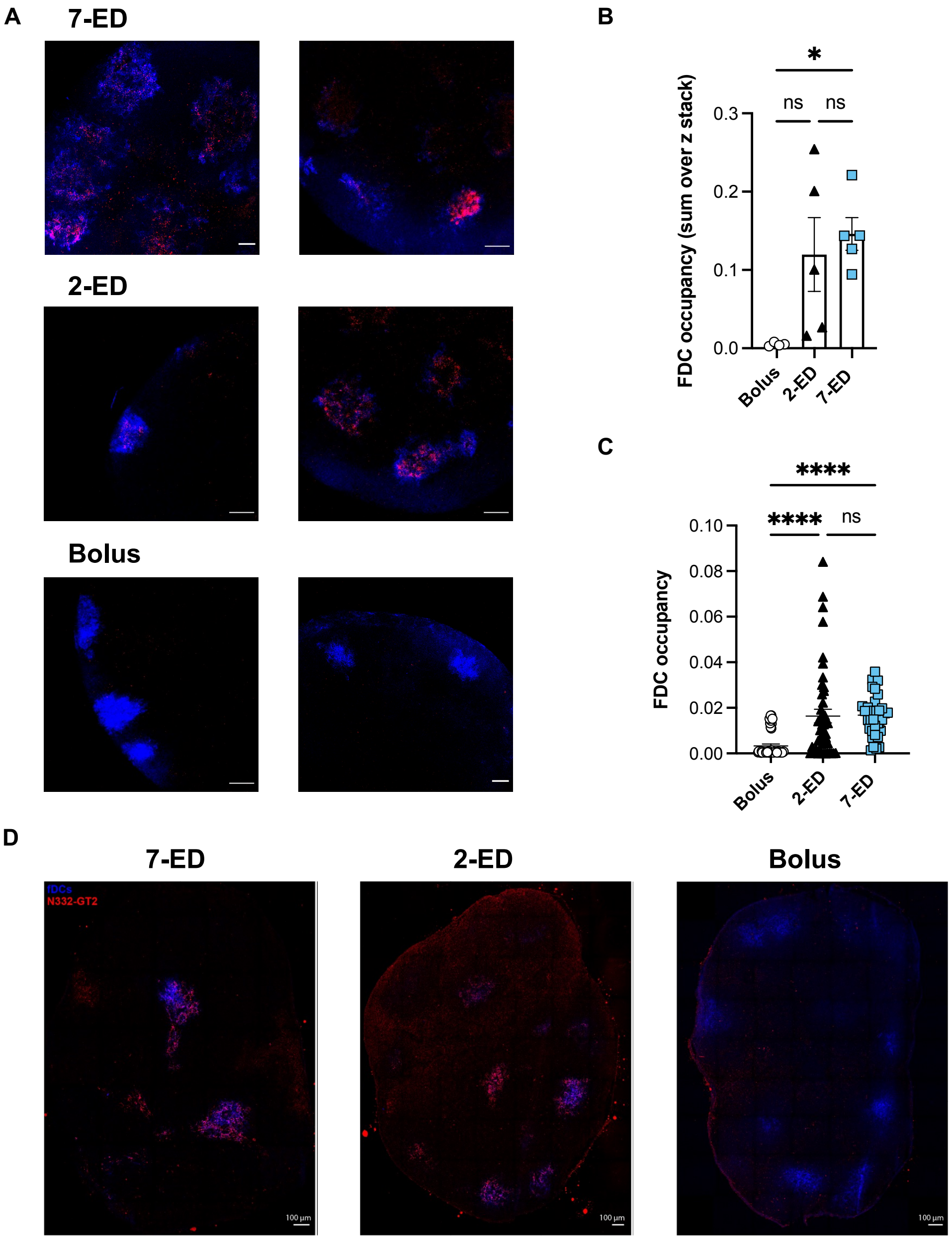


**figure S4. Additional antigen capture on FDCs data. (A-D)** Groups of C57Bl/6 mice (*n* = 3 animals/group) were immunized by bolus, 2-ED, or 7-ED regimens as shown in Fig. 3G, followed by collection of lymph nodes for imaging at 48 h after bolus or after the last injection of the 2-ED and 7-ED regimens. Shown are additional cleared whole LN images for bolus, 2-ED, and 7-ED regimens **(A)**, fraction of FDC area occupied by antigen for each individual image in the z-stack (9 per image) **(B)**, and for the z-projection (sum of slices) **(C)** and additional representative histological LN slices for 7-ED, 2-ED, and bolus dosing regimens **(D)**. Shown are data from one independent experiment for each immunization series. ****, $p < 0.0001$; ***, $p < 0.001$; **, $p < 0.01$; *, $p < 0.05$; ns, not significant; by one-way ANOVA with Dunnett's multiple comparison post test compared to bolus immunization.

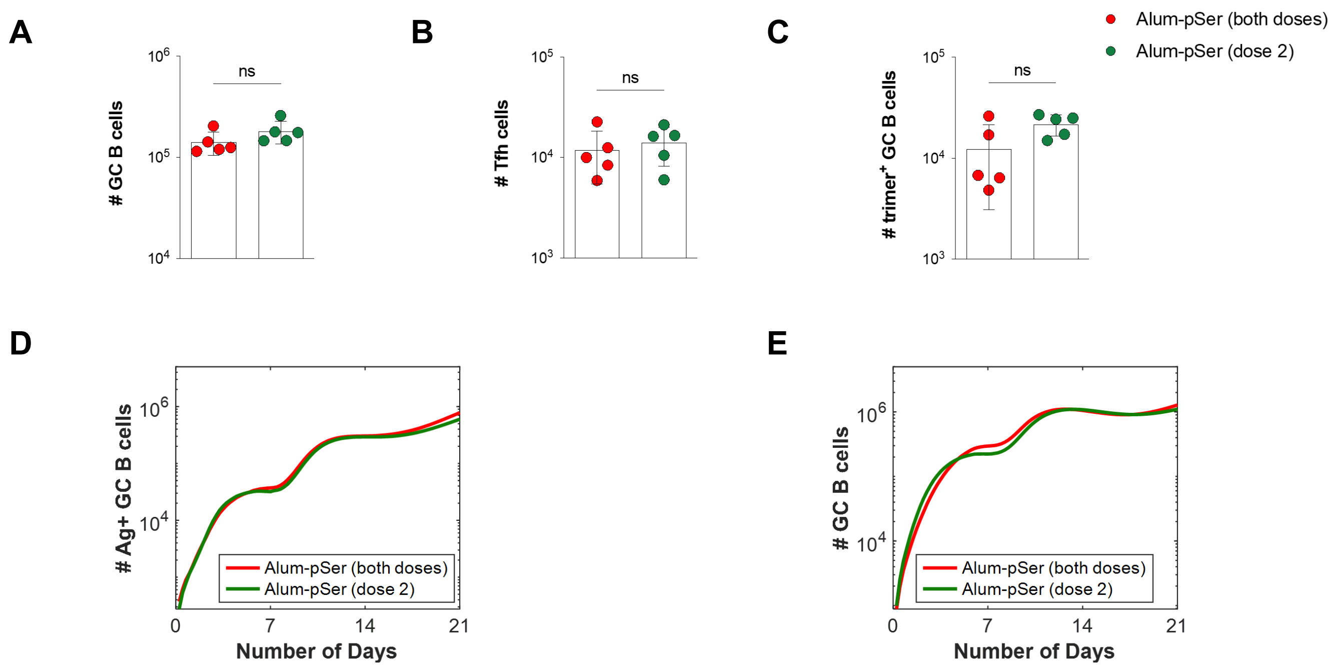


**figure S5: Additional data on Alum-pSer immunizations. (A-C)** C57Bl/6J mice (*n*=5 animals/group) were immunized with 10 µg MD39 trimer anchored onto 50 ug Alum combined with 5 µg SMNP adjuvant (20% vaccine on day 0, 80% on day 7, red) or with alum anchoring only used for the second dose (green). Shown are experimentally measured responses on day 14 for GC B cells **(A),** Tfh cells **(B),** and trimer-specific GC B cells **(C). (D-E)** *In silico* predictions modeling the same experiment as in (A-C), showing numbers of native antigen-binding GC B cells **(D),** and total GC B cells **(E)** over time. Shown are data from one independent experiment for each immunization series. ****, $p < 0.0001$; ***, $p < 0.001$; **, $p < 0.01$; *, $p < 0.05$; ns, not significant; by one-way ANOVA with Dunnett's multiple comparison post test compared to bolus immunization.

**Table S1. Antigen decay kinetics data.**

Measured antigen trimer mass per lymph node after immunization. Maximum likelihood estimate of the antigen decay rate assuming first-order kinetics is ~2.7 / day, corresponding to half-life of ~6.2 hrs.

| | **Extracellular Ag (ng)** | | |
|---|---|---|---|
| **6 Hrs** | 52.11 | 38.29 | 48.32 |
| **Day 1** | 5.60 | 6.85 | 6.82 |
| **Day 2** | 0.64 | 0.12 | 0.83 |

### Table S2. Simulation parameters for the model of T cell priming

Descriptions and values of the parameters used in the simulation of T cell priming.

| Parameter | Value | Description | Equation | Note |
|---|---|---|---|---|
| **T cell priming** | | | | |
| $d_{Ag}$ | $3\ \text{day}^{-1}$ | Antigen decay rate | 1 | Our data (Table S1) |
| $d_{Adj}$ | $3\ \text{day}^{-1}$ | Adjuvant decay rate | 2 | Taken to be identical to $d_{Ag}$ |
| $S_{Adj}$ | 0.1 | Half-max adjuvant activity concentration | 3 | See text for Eq. 3 |
| $k$ | 10 | Max increase in antigen uptake rate | 4, 5 | See text for Eqs. 4-5 |
| $\mu$ | 1.2 | Innate immune cell death rate | 3, 4, 5 | From ref. (*37*) |
| $\alpha$ | 1.5 | Max T cell proliferation rate | 6 | |
| $\eta$ | 0.22 | T cell differentiation rate | 7 | |
| $D_0$ | $1.8 \times 10^6$ | Rate of DC recruitment | 4 | Fitted to data (Fig. 3F) |
| $T_0$ | 28 | Baseline number of T cells | 6,7 | Fitted to data (Fig. 3F) |
| $[Ag]_0$, $[Adj]_0$, $[TC]_0$, $[DC]_0$, $[aDC^{Ag+}]_0$, $[T_{FH}]_0$ | 0 | Initial conditions | 1-7 | |
| $[T]_0$ | $T_0$ | | | |

**Table S3. Simulation parameters for the model of B cell responses**

Descriptions and values of the parameters used in the simulation of B cell responses. Highlighted parameters have values changed from the model in ref. (*35*).

| Parameter | Value | Description | Equation | Note |
|---|---|---|---|---|
| **Antigen and antibody dynamics** | | | | |
| $k_{Ig}$ | $10^{-1}$ nM day$^{-1}$ PC$^{-1}$ | Rate of antibody production per plasma cell per day, in terms of change in serum concentration | 13, 14 | Increased from ref. (*35*) to match the faster kinetics observed in mice |
| $d_{Ig}$ | 0.17 day$^{-1}$ | Antibody decay rate | 13 | |
| $d_{Ag}$ | 3 day$^{-1}$ | Antigen decay rate | 9,10 | From ref. (*35*) |
| $k_{deposit}$ | 1 hour$^{-1}$ | Rate of immune complex transport to FDC | 11, 12 | |
| $d_{IC}$ | 0.15 day$^{-1}$ | Rate of decay of immune complex on FDC | 12 | |
| $[Ag]_0$ | 0 nM | Initial Conditions | 8-14 | |
| $[Ig]_0$ | $10^{-2}$ nM | | | |
| $[IC]_0$, $[IC-FDC]_0$ | 0 nM | | | |
| **B cell affinities** | | | | |
| $N_{naive}$ | 2000 cells/GC | Number of naïve B cells / GC | 16, 17 | From ref. (*35*) |
| $p$ | 0.5 | Fraction of naïve B cells that target the naitve antigen | 18, 19 | |
| $E_1^h$ | 7.2 | Affinity at which there is one dominant naive B cell available for each GC on average | 18, 19 | |
| $dE_{12}$ | 0.6 | $E_1^h - dE_{12}$ is the affinity at which there is one subdominant naïve B cell available for each GC on average | 18, 19 | |
| $n_{res}$ | 80 | Length of the string representation of B cell residues | 24 | |
| $\mu, \sigma, \epsilon$ | 3.1, 1.2, 3.08 | Parameters for the shifted log-normal distribution that represent the effects of mutations on B cell binding affinities | 25 | |
| **GC and EGC** | | | | |
| $C_0$ | 0.2 nM | Reference antigen concentration | 20 | Increased from ref. (*35*) to match the faster kinetics observed in mice |
| $\beta_{max}$ | 4 day$^{-1}$ | Maximum rate of positive selection for GC and EGC B cells | 23 | |
| $E_0$ | 6 | Reference binding affinity | 20 | From ref. (*35*) |
| $K$ | 0.5 | Stringency of selection of naïve and GC B cells by helper T cells based on the amounts of antigen internalized | 20 | |
| $N_{T0}$ | 1200 | Maximum number of helper T cells | 15 | |
| $\alpha$ | 0.5 day$^{-1}$ | Death rate of GC B cells | 6 | |

| | | | | |
|---|---|---|---|---|
| **Memory and Plasma Cell Dynamics** | | | | |
| $p_1$ | 0.1 | Probability that a positively selected GC B cell exits by differentiation | Text | From ref. (*35*) |
| $p_2$ | 0.1 | Probability that a differentiating GC B cell becomes a plasma cell | | |
| $p_2^{EGC}$ | 0.6 | Probability that a proliferating memory cell in EGC differentiates into a plasma cell | | |
| $d_{PC}$ | 0.17 $\text{day}^{-1}$ | Death rate of plasma cells | | |

### Details of Computational Models

#### Model of T cell priming

We constructed our model based on the biological observations of the mechanism of action of adjuvants (*55-60*). Upon administration of adjuvants, local tissue-resident cells such as neutrophils and macrophages are recruited to the site of vaccine administration and draining lymph nodes. These cells release cytokines and chemokines, which serve as chemotactic agents for dendritic cells (DCs). As a result, DCs migrate to the sites and take up antigen. The presence of adjuvant significantly enhances this process. Adjuvant facilitates the maturation of DCs, ensuring a more efficient antigen uptake (*59*), and aids in the delivery of antigen to the DCs (*30*). Following antigen uptake, the activated DCs present peptide-major histocompatibility complex (pMHC) molecules to CD4 T cells in lymph nodes. This interaction leads to the proliferation of CD4 T cells and initiates their differentiation into Tfh cells (*44*).

In our model, the concentrations of antigen and adjuvant rise upon administration and subsequently decay according to first-order kinetics, consistent with previous models (*19, 35, 37*). Regardless of the dosing scheme, the same total quantity of antigen is administered. Thus, we normalize the concentrations so that the total amount correspond to a concentration of 1. The same normalization is applied to the adjuvant concentration. For a dosing scheme involving *n* doses given at times $t_1, \dots t_n$, the changes in concentration of the antigen from each dose are represented by $f_{1,Ag}, \dots f_{n,Ag}$, and of adjuvant are represented $f_{1,Adj}, \dots f_{n,Adj}$. These fractions satisfy the conditions $\sum_{i=1}^{n} f_{i,Ag} = 1$ and $\sum_{i=1}^{n} f_{i,Adj} = 1$, respectively. Then, the differential equations that govern the antigen and adjuvant concentrations are as follows:

$$\frac{d[Ag]}{dt} = \sum_{i=1}^{n} f_{i,Ag}\delta(t - t_i) - d_{Ag}[Ag] \qquad \text{(Equation 1)}$$

$$\frac{d[Adj]}{dt} = \sum_{i=1}^{n} f_{i,Adj}\delta(t - t_i) - d_{Adj}[Adj] \qquad \text{(Equation 2)}$$

where $\delta(t - t_i)$ is the Dirac delta function, whose value is zero everywhere except at $t = t_i$, and whose integral over the domain that includes $t = t_i$ is equal to one.

The recruitment of tissue-resident innate immune cells (TCs) and their decay are modeled as:

$$\frac{d[TC]}{dt} = \frac{[Adj]}{S_{Adj} + [Adj]} - \mu[TC] \qquad \text{(Equation 3)}$$

where the first term stands for the activity of the adjuvant with $S_{Adj}$ being the half-max adjuvant concentration. The activity increases with adjuvant concentration when $S_{Adj} > [Adj]$ but saturate when $S_{Adj} \ll [Adj]$. Employing a saturation function to represent biological activity is widely adopted (*35*, *37*, *61*). We pick a small value for $S_{Adj}$ based on the observation that there is not a big difference between the numbers of DCs recruited after full dose and 20 % of the dose (Fig. S3H). $\mu$ is the decay rate, assumed to be identical for all innate immune cells for simplicity. Its value is taken from Mayer et al. (*37*) The recruitment of DCs by the TCs and their activation and antigen uptake are modeled as:

$$\frac{d[DC]}{dt} = D_0[TC] - \left(1 + k\frac{[Adj]}{S_{Adj} + [Adj]}\right)[DC][Ag] - \mu[DC] \qquad \text{(Equation 4)}$$

$$\frac{d\left[aDC^{Ag^+}\right]}{dt} = \left(1 + k\frac{[Adj]}{S_{Adj} + [Adj]}\right)[DC][Ag] - \mu\left[aDC^{Ag^+}\right] \qquad \text{(Equation 5)}$$

where $aDC^{Ag^+}$ represent activated antigen-loaded DCs. Here, $D_0$ is the rate of DC recruitment and its value is derived from the best fit of model prediction to experimental data in Fig. 3F. The parameter $k$ quantifies the extent to which the adjuvant's activity expedite antigen uptake by the DCs. The adjuvant, SMNP, is a saponin/Toll-like receptor 4 agonist that can activate DCs and increase their antigen uptake (*30*). Given this recognized role of adjuvants, we set $k$ to be large.

The activated antigen-primed DCs induce the proliferation of CD4 T cells, which we assume to differentiate into Tfh cells according to first-order kinetics:

$$\frac{d[T]}{dt} = \alpha\frac{[aDC^{Ag^+}][T]}{[T] + [aDC^{Ag^+}]} - \eta([T] - T_0) \qquad \text{(Equation 6)}$$

$$\frac{d[T_{FH}]}{dt} = \eta([T] - T_0) \qquad \text{(Equation 7)}$$

where $\alpha$ is the maximum proliferation of the T cells, $\eta$ is the rate at which proliferating T cells differentiate into Tfh cells, and $T_0$ is the baseline number of T cells. The model of T cell proliferation and its parameter values for $\alpha$ and $\eta$ are derived from Mayer et al. (*37*), which encompasses T cell proliferation and death. In this study, rather than accounting for death, we model the process as T cells proliferating and differentiating (*62*).

We fit two parameters, $D_0$ and $T_0$, based on finding the best alignment between the model prediction and experimental data of Tfh cell numbers in Fig. 3F. We utilized standard maximum likelihood estimation method. $D_0$ controls the scale of the number of DCs and $aDC^{Ag+}$s, while $T_0$ controls the scale of the number of T cells. Since the ratio between $aDC^{Ag+}$s and T cells affect the proliferation rate of T cells (Eq. 6), appropriate scaling between $D_0$ and $T_0$ is required to recapitulate the fold-difference between the experimentally observed Tfh numbers after bolus and the ED schemes. If the scale of DCs is too small compared to the scale of baseline T cells, the initial doses containing small amounts of antigen and adjuvant will not generate enough $aDC^{Ag+}$s for adequate T cell proliferation. Thus, fold-difference between bolus and the ED schemes is reduced. When the scale of DCs is too large, the opposite is true. Additionally, the absolute value of $T_0$ affects the overall quantitative match between the experimental and simulated data. The antigen decay rate was approximated from experimental data in Fig. S1. All other parameters were derived from previous literature or reasonable assumption. See Table S2 for the summary of model parameters and their values.

### Model of B cell response

We adapted and refined the B cell response model from our previously published study (*35*), where an in-depth description of the rationale for model development, alternative model structure exploration, and parameter sensitivity analysis can be found. Here, we present a concise overview of the model as applied in our study, including the equations and parameters necessary for reproducing the results. Additionally, we have outlined the changes made to the original model to aid readers interested in understanding the modifications. See Table S3 for a summary of parameters and their values.

#### *Overview of the model*

The model incorporates four key components of B cell and antibody dynamics: (i) display of antigens on follicular dendritic cells (FDCs), (ii) activation of naïve B cells and entrance into GCs, (iii) the process of affinity maturation in GCs, leading to the production of memory and plasma cells, and (iv) the process of memory B cell expansion and differentiation outside of GCs upon antigen encounter. To capture the dynamics of antigen administration, breakdown, immune complex formation, and transport to FDCs, we employ a system of differential equations. Immune complex formation depends on the concentration and affinity of antigen-specific antibodies, which is determined by the model for B cell development and antibody production.

For simulating the various stages of B cell activation, proliferation, mutation, and differentiation, we utilize a stochastic agent-based model. We categorize B cells into four classes: naive B cells, GC B cells, memory B cells, and plasma cells. The simulation progresses in time steps of 0.01 day, and antigen and antibody concentrations are updated at each step. Simultaneously, in each step, probabilities for events like activation, selection, cell division, somatic hypermutation, differentiation, and apoptosis are computed for the B cells and subsequently executed. To emulate the conditions of a secondary lymphoid organ, the simulation simultaneously models up to 200 distinct GCs. The number of Tfh cells obtained from the model of T cell priming is used to determine the number of active GCs and the number of Tfh cells available in each GC. Each simulation is repeated 10 times. The averaged outcomes can be interpreted as representative of a population-level immune response.

In this study, we consider the partial degradation of antigen as schematically depicted in Fig. 4A. We postulate there are two distinct antibodies: one targeting the native antigen (nAg) and the other targeting the partially degraded, non-native antigen (pAg). We further postulate that the partially degraded antigen is immunodominant. Drawing parallel to our original model - which distinguished immunodominant and subdominant epitopes on the SARS-CoV-2 receptor binding site (*35*), we employ a two-epitope model. In this approach, epitopes on the nAg are aggregated into a single subdominant epitope while the epitopes on the pAg are aggregated into a single immunodominant epitope. Each B cell targets either the nAg or the pAg. In this model, the germline affinity distribution of naïve B cells displays an extended tail for the immunodominant non-native antigen.

#### *Equations for the antigen dynamics*

First, we describe the processes that are included in the model for antigen dynamics. We use the following abbreviations: native soluble antigen (nAg), partially degraded soluble antigen (pAg), either type of soluble antigen (Ag), soluble antibody (Ig), soluble immune complex (IC), immune complex on follicular dendritic cell (IC-FDC), plasma cell (PC).

The partial and full decay of the soluble antigen are described as:

| | |
|---|---|
| $nAg \rightarrow pAg$ | Decay of free native soluble antigen into partially degraded antigen |
| $pAg \rightarrow \emptyset$ | Decay of partially degraded antigen |

The production and decay of antibodies, which apply to both the nAg and pAg-targeting antibodies are described as:

| | |
|---|---|
| $PC \rightarrow PC + Ig$ | Antibody production by plasma cells |
| $Ig \rightarrow \emptyset$ | Decay of free soluble antibody |

The formation of immune complexes, deposition on FDC, and decay on FDC, which apply to both the nAg and pAg, are described as:

| | |
|---|---|
| $Ag + Ig \rightleftharpoons IC$ | Fast equilibrium between free soluble antigen and antibody |
| $IC \rightarrow IC - FDC$ | Immune complex transport to follicular dendritic cells |
| $IC - FDC \rightarrow \emptyset$ | Consumption and decay of immune complexes on follicular dendritic cells |

The above processes lead to the following differential-algebraic equations:

$$\frac{[Ag][Ig]}{[IC]} = K_d \quad \text{(Equation 8)}$$

$$\frac{d[nAg]}{dt} = -d_{Ag}[Ag] \quad \text{(Equation 9)}$$

$$\frac{d[pAg]}{dt} = d_{Ag}[nAg] - d_{Ag}[pAg] \quad \text{(Equation 10)}$$

$$\frac{d[IC]}{dt} = -k_{deposit}[IC] \quad \text{(Equation 11)}$$

$$\frac{d[IC - FDC]}{dt} = k_{deposit}[IC] - d_{IC}[IC - FDC] \quad \text{(Equation 12)}$$

$$\frac{d[Ig]}{dt} = k_{Ig}[PC] - d_{Ig}[Ig] \quad \text{(Equation 13)}$$

$$\frac{dK_d}{dt} = \frac{(K_d^{PC} - K_d)k_{Ig}[PC]}{[Ig] + [IC]} \quad \text{(Equation 14)}$$

Here, $K_d$ and $K_d^{PC}$ represent the dissociation constants of the serum antibodies and the BCRs of plasma cells, respectively.

#### ***Equations for the B cell dynamics***

<u>*Initiation of GCs*</u>

In Yang et al. (*35*), which solely focused on bolus injections, the assumption was made that 200 GCs are simultaneously initiated post each injection, with a separate pool of naïve B cells tied to each GC. This simplified model is not best suited for depicting dosing schemes with gradual administration of antigen and adjuvant. Here, we consider a single pool of naïve B cells collectively shared across all GCs. We then postulate that GCs are sequentially initiated, influenced by the increasing number of Tfh cells. The number of Tfh cells is determined by the model of T cell priming (Eqs. 1-7) and is affected by the dosing scheme.

The number of active GCs at time *t*, $N_{GC}(t)$, is determined from the number of Tfh cells, $N_{Tfh}(t)$, as follows:

$$N_{GC}(t) = \min(\left\lceil \frac{N_{Tfh}(t)}{N_{T0}}, 200 \right\rceil) \qquad \text{(Equation 15)}$$

While the total number of GCs has not reached the capacity of 200, $N_{T0}$ quantifies the highest number of Tfh cells that can be attributed to any one GC. As Tfh cell numbers rise, new GCs are initiated. However, once the capacity of 200 GCs is reached, further increase in the number of Tfh cells does not lead to establishment of new GCs. Instead, these additional Tfh cells are distributed among the existing GCs, resulting in larger individual GCs. From our simulations, bolus immunization leads to an average of ~60 GCs, while 2-ED results in ~187 GCs, and 7-ED reaches the maximum capacity with 200 GCs. In terms of the average number of B cells per GC, both bolus and 2-ED present ~1400 B cells, whereas 7-ED displays significant increase to ~5900 B cells.

<u>*Naïve B cell germline affinities*</u>

The model of germline-endowed binding affinities of naïve B cells follows the approach from the previous study We parametrize the distribution of the binding affinity, denoted as $E = -\log_{10}(K_d)$, with a geometric distribution. The distribution takes discrete values between 6 and 8, expressed as $E_k = 6 + 0.2k$ for $k = 0 \dots 10$. Note that a higher value of $E$ indicates stronger binding.

We differentiate between two groups of naïve B cells: one group binding to the native antigen and the other to the partially degraded antigen. We assume that the partially degraded antigen is more immunodominant. The frequencies of naïve B cells binding to either antigen type with affinity $E_k$ are formulated as follows:

$$f_{pAg}(E_k) = N_{naive}(1-p)\frac{e^{-r_1 E_k}}{\sum_{k=0}^{10} e^{-r_1 E_k}} \quad \text{(Equation 16)}$$

$$f_{nAg}(E_k) = N_{naive}p\frac{e^{-r_2 E_k}}{\sum_{k=0}^{10} e^{-r_2 E_k}} \quad \text{(Equation 17)}$$

where $N_{naive}$ is the number of naïve B cells and $p$ is the fraction of B cells that target the native antigen. The exponents $r_1$, $r_2$ determine the slope of the distribution. They are derived from the parameters $E_1^h$ and $dE_{12}$ which we directly specify. The following relationships are satisfied:

$$f_{pAg}\left(E_1^h\right) = 1-p \quad \text{(Equation 18)}$$

$$f_{nAg}\left(E_1^h - dE_{12}\right) = p \quad \text{(Equation 19)}$$

Given $dE_{12} > 1$, this parameterization results in an extended tail of germline-endowed affinities for the naïve B cells targeting the partially degraded antigen, reflecting their immunodominance.
The above model was initially developed to represent a pool of naïve B cells associated with an individual GC. For this study, we adjust the model by amplifying the frequencies by a factor of 200, thereby consolidating a shared pool of naïve B cells across all GCs.

_Activation of naïve B cells, differentiation into plasmablasts, and entry into GCs_

We model the quantity of antigen captured by a B cell, denoted as $i,$ using the following equation:

$$A_i = \left(\frac{C}{C_0}10^{(\min(E_i,10)-E_0)}\right)^K \quad \text{(Equation 20)}$$

where $\frac{C}{C_0}$ signifies antigen availability, $E_0 = 6$ serves as the reference affinity, and $K$ represents the selection stringency. The *min* function accounts for the effect of affinity ceiling. The term $C = 0.01([Ag] + [IC]) + [IC - FDC]$ represents the effective antigen concentration, accounting for the predominant influence of surface-presented immune complexes. $C_0$ is the reference concentration. For B cells targeting the native or partially degraded antigen, the appropriate antigen concentration is applied respectively.
The probability of B cell activation at each step is determined as:

$$\Pr(\text{B cell } i \text{ is activated}) = \min(A_i, 1) \quad \text{(Equation 21)}$$

The entry of activated naïve B cells to GCs is limited by competition for positive selection from helper T cells. The selection rate for an activated naive B cell $i$, denoted as $\lambda_i$, is given by:

$$\lambda_i = \frac{\frac{N_T}{N_{activated}} \frac{A_i}{\langle A \rangle}}{1 + \frac{N_T}{N_{activated}} \frac{A_i}{\langle A \rangle}} \quad \text{(Equation 22)}$$

Here, $N_{activated}$ stands for the total count of activated B cells, $\langle A \rangle$ denotes the average quantity of antigen captured by all activated B cells, and $N_T$ represents the limited number of helper T cells that B cells compete for. Activated naïve B cells migrate to the T-B border and interact with antigen-primed T cells before they migrate back to the B cell zone and enter GCs (*62*). Thus, we choose the value of $N_T$ as the number of T cells from the model of T cell priming (Eq. 6).

Upon positive selection, a naïve B cell has three potential fates: it can differentiate into a memory cell, a plasmablast, or a GC B cell (*63*). In our model, there is a probability $p_1$ that a B cell differentiates into either a memory cell or a plasmablast. If it takes this path, it further has a probability $p_2$ of becoming a plasma cell and otherwise becomes a memory cell. If the B cell becomes a plasmablast, it undergoes five division cycles (*64*). Conversely, there is a probability 1- $p_1$ that the B cell enters a GC. If this path is taken, a naïve B cell is randomly allocated to one of the active GCs.

*Proliferation, mutation, and death of GC B cells*

GC B cells must be activated by antigen and then receive help from Tfh cells to proliferate. The activation step for GC B cells is identical to that of naïve B cells. The rate of positive selection, $\beta_i$, of a GC B cell, $i$, is modeled based on competition for limited number of Tfh cells:

$$\beta_i = \beta_{max} \frac{\frac{N_{Tfh}/N_{GC}}{N_{activated}} \frac{A_i}{\langle A \rangle}}{1 + \frac{N_{Tfh}/N_{GC}}{N_{activated}} \frac{A_i}{\langle A \rangle}} \quad \text{(Equation 23)}$$

where $\beta_{max}$ is the maximum rate of positive selection, and the number of Tfh cells in each active GC is calculated as $N_{Tfh}/N_{GC}$.

Upon positive selection, a B cell exits the GC with a probability of $p_1$ and proliferates inside the GC with a probability of 1-$p_1$. When it exits the GC, it becomes a plasma cell with a probability of $p_2$ or a memory cell with a probability of 1- $p_2$. When it proliferates, one offspring undergoes mutation. This mutation can result in apoptosis (probability 0.3), no affinity change (probability 0.5), or a change in the mutation state of a randomly selected residue (probability 0.2).

The affinities of GC B cells change with mutations. Each B cell is represented as a string of 0's and 1's with a total length of $n_{res}$. The residues are all 0's for a naïve B cell. Each time an affinity-affecting mutation occurs, one of the residues is randomly chosen and its value is flipped. The affinity of a B cell is determined by the sum of its germline-endowed affinity and the contributions of the mutations, as follows:

$$E_i = E_i^{germline} + \sum_{j=1}^{n_{res}} \delta_{i,j} s_{i,j} \qquad \text{(Equation 24)}$$

where $\delta_{i,j} \in \{0,1\}$ is the mutational state of residue $j$, and $s_{i,j}$ is the effect of the mutation at residue $j$ on the binding affinity. The values of $s_{i,j}$ $(j = 1 \dots n_{res})$ are independent and identically distributed as follows:

$$s_{i,j} \sim e^{N(\mu,\sigma)} - \epsilon \qquad \text{(Equation 25)}$$

This shifted log-normal distribution with $\mu, \sigma, \epsilon$ chosen to fit experimentally determined distribution leads to ~5 % of affinity-affecting mutations increasing the binding affinity.

<u>*Expansion and differentiation of B cells in Extra Germinal Center*</u>

In schemes involving multiple injections, we model the expansion of memory cells and their differentiation into plasma cells outside of germinal centers, termed extra germinal centers (EGCs). Any injection beyond the first dose initiates an EGC, unless there is already an ongoing one. EGC proliferation is rapid and short-lived (*65*). An EGC terminates after a 6-day period without a new injection. The process of affinity-based positive selection, which results in either differentiation or proliferation, is identical to that in the GCs. However, no mutations are introduced to B cells in EGCs, and proliferating cells in EGC differentiates into plasma cells with higher probability, $p_2^{EGC}$ (*65*, *66*). Moreover, to reflect the fast kinetics in the EGC, the number of Tfh cells is maintained at its peak value, $N_{T0}$.

***Modifications from the original model***

In our study, we have introduced several changes from the original model proposed by Yang et al. (*35*), which are described in detail in the relevant sections. We summarize these changes here to aid readers interested in understanding the modifications.

(1) T cell priming: We developed a detailed T cell priming model that is suitable for gradual dosing schemes.
(2) Antigen decay and targeting: We introduced partial decay of antigen and B cells targeting the partially degraded antigen.
(3) Initiation of GCs: We introduced asynchronous initiation of GCs, which is suitable for gradual dosing schemes.

(4) Cell differentiation: We consider differentiation of memory cells and plasmablasts from a positively selected naïve B cell. This was not considered for simplicity in the previous model, but the shorter timescale in this study makes these early dynamics important.

(5) Parameter adjustment: We adjusted a small number of model parameters to match the timescales of experimental observations in the current study. The previous model was built for vaccine responses in humans, which have slower and longer-lasting GC responses than mice. Table S3 provides all the parameter values for the model of B cell responses. Those that take values not tested in Yang et al. are highlighted.
